# The columnar structure of human telomeric chromatin suggests mechanisms for telomere maintenance

**DOI:** 10.1101/2022.06.15.496090

**Authors:** Aghil Soman, Sook Yi Wong, Nikolay Korolev, Wahyu Surya, Simon Lattman, Vinod K. Vogirala, Qinming Chen, Nikolay V. Berezhnoy, John van Noort, Daniela Rhodes, Lars Nordenskiöld

## Abstract

Telomeres, the ends of eukaryotic chromosomes, play pivotal roles in ageing and cancer and are targets of DNA damage and response. However, little is known about the structure and organization of telomeric chromatin at the molecular level. We used electron microscopy and single-molecule magnetic tweezers to characterize well-defined telomeric chromatin fibers of kilobasepair length. The cryo-EM structure of the compact telomeric tetranucleosome revealed a novel columnar folding, unusually short nucleosome repeat length of ∼132bp and the role of the histone N-terminal tails in stabilizing this structure. This is the first near-high resolution structure of chromatin with a native DNA sequence. The columnar structure exposes the DNA, making them susceptible to DNA damage. The telomeric tetranucleosome also exists in an alternative well-defined state, with one nucleosome open, accessible to protein factors. This suggests that protein factors, which plays a role in maintaining telomeres, can bind to telomeric chromatin in its compact heterochromatic form. The features of the telomeric chromatin structure reveals important insights of significant relevance for telomere function *in vivo* that provides information on mechanisms of nucleosome recognition by chromatin factors

---

Mammalian telomeres consist of the conserved and tandemly arranged DNA sequence repeat TTAGGG^1^. Telomeric repeats act as a platform for recruiting telomere-specific binding factors, forming the capping structure at the end of linear eukaryotic chromosomes that protect chromosome ends from incorrect DNA repair and degradation^2^. This capping structure consists of the Shelterin complex (a complex of six different proteins) and nucleosomes, packaging 5-15 kbp of double-stranded telomeric DNA in humans^1–4^. Aberrant regulation of telomeres has been linked with tumour development and pathologies of ageing. However, the detailed structural information on the interaction of shelterin and its components with telomeric DNA and nucleosomes is unknown. Furthermore, despite their essential role in protection, telomeres are hotspots for DNA damage, and dysfunctional telomeres are prime targets of DNA damage response (DDR)^5–8^. The reasons for this are not well understood.

The DNA of eukaryotic genomes is packaged into chromatin, comprising a linear array of nucleosome core particles (NCP) consisting of about 147 bp of DNA wrapped around a histone octamer (HO) composed of two copies each of the four histone proteins, H2B, H3, and H4. Genome-wide nucleosome positioning studies revealed an average nucleosome repeat length (NRL) that ranges from 167 bp in *S. cerevisiae* to 197 bp in mammalian cells^9, 10^. Depending on the NRL and the presence of cations and bound linker histones, nucleosome arrays fold into higher-order chromatin structures of different topologies and compaction, including the so-called 30-nm fibres^11^. In recent years, pioneering EM, X-ray crystallography, and force spectroscopy studies employing the Widom ‘601’ nucleosome positioning DNA sequence have provided unprecedented insights into the structure and folding dynamics of chromatin fibres^12–16^.

Telomeric DNA is known to form chromatin, and nuclease digestion studies revealed that its NRL appears exceptionally short – about 157 bp with a broad distribution of lengths in the range of ±30 bp^17, 18^. Our recently determined crystal structure of the telomeric NCP at 2.2 Å resolution^19^ revealed a three-dimensional structure similar to other NCPs^20, 21^. However, we also found that it displayed a less stable and markedly more dynamic nucleosome than NCPs containing DNA positioning sequences. The explanation for this is based on the physical properties of the G-rich telomeric TTAGGG six bp repeat, disfavoring nucleosome positioning and rendering telomeric nucleosomes more mobile^22^. How the short NRL and this lower stability of telomeric nucleosomes affect the structure and dynamics of telomeric chromatin at the molecular level is unknown.

To fill this knowledge gap, we produced telomeric nucleosome arrays of length representative of natural telomeres and characterised their properties using a combination of electron microscopy and single-molecule magnetic tweezers. We found that kbp-long fibres at extended beads-on-a-string conditions showed heterogeneous nucleosome positioning and that arrays folded in the presence of Mg^2+^ displayed distinct columnar packing. Structural analysis of the folded telomeric tetra-nucleosome revealed a novel columnar structure with close nucleosome stacking. The structure of the di-nucleosome (Di-NCP) unit at 3.9 Å resolution exhibited close stacking of NCPs with an unusually short NRL of around 132 bp with DNA wound in a continuous superhelix around the stacked HOs. Furthermore, tail-mediated interactions between nucleosomes by DNA-DNA bridging was revealed. In addition, we identified a distinct ‘open’ conformation where one NCP is flipped out from its stacked position, with the two NCP disks perpendicular to each other and accessible to chromatin modifiers. To the best of our knowledge, this is the first near-the high-resolution structure of chromatin on a natural DNA sequence and the first telomeric chromatin structure.

## Telomeric DNA array produced by large-scale ligation

We developed a general and flexible method that enabled the preparation of large quantities of telomeric DNA with a specific number of telomeric repeats (Methods and Extended Data Fig. 1a-c). In brief, 157 bp telomeric DNA consisting entirely of TTAGGG repeats flanked by AvaI restriction sites was multimerized to produce a DNA array containing up to 20×157 bp telomeric DNA repeats (Telo-20 DNA) through the *in-vitro* ligation of two Telo-10 constructs (Extended Data Fig. 1d). For the EM and magnetic tweezers experiments, a 20-mer hybrid construct with 18 telomeric repeats flanked by 157 bp of ‘601’ nucleosome positioning DNA sequence was also produced (#Telo-18# DNA) (Extended Data Fig. 2a and Extended Data Fig. 2d). In the single-particle Cryo-EM analysis, DNA array templates comprising 4×157 bp telomeric DNA (Telo-4 DNA) and 4×157 bp Widom 601 DNA were used (Extended Data Fig. 2f). The DNA array templates were reconstituted with a recombinant human HO (Extended Data Fig. 2g-l)^23^.

## Telomeric nucleosome arrays show beads-on-a-string conformation with heterogeneous nucleosome distribution at low salt conditions

Reconstituted Telo-20 and hybrid #Telo-18# nucleosome arrays (the latter, which can also be characterised by force spectroscopy, see below) were visualized using negative stain EM. At low salt conditions, these arrays displayed the characteristic ‘beads-on-a-string” conformation (Fig. 1a-c). The Telo-20 fibres showed a broad nucleosome distribution ranging from 14-23 nucleosomes per array (median 17) and variable nucleosome spacing (Fig. 1a and d). The #Telo-18# nucleosome arrays showed a more homogeneous occupancy in the range of 16-20 nucleosomes per array (median 18) (Fig. 1b and d). In both telomeric arrays, we observed clusters of tightly packed nucleosomes and regions with nucleosomes separated by larger distances (Fig. 1a and b). As expected, the 601-20 control arrays showed evenly distributed nucleosomes containing 18-20 nucleosomes (median 20) (Fig. 1c and d).

**Fig. 1.**
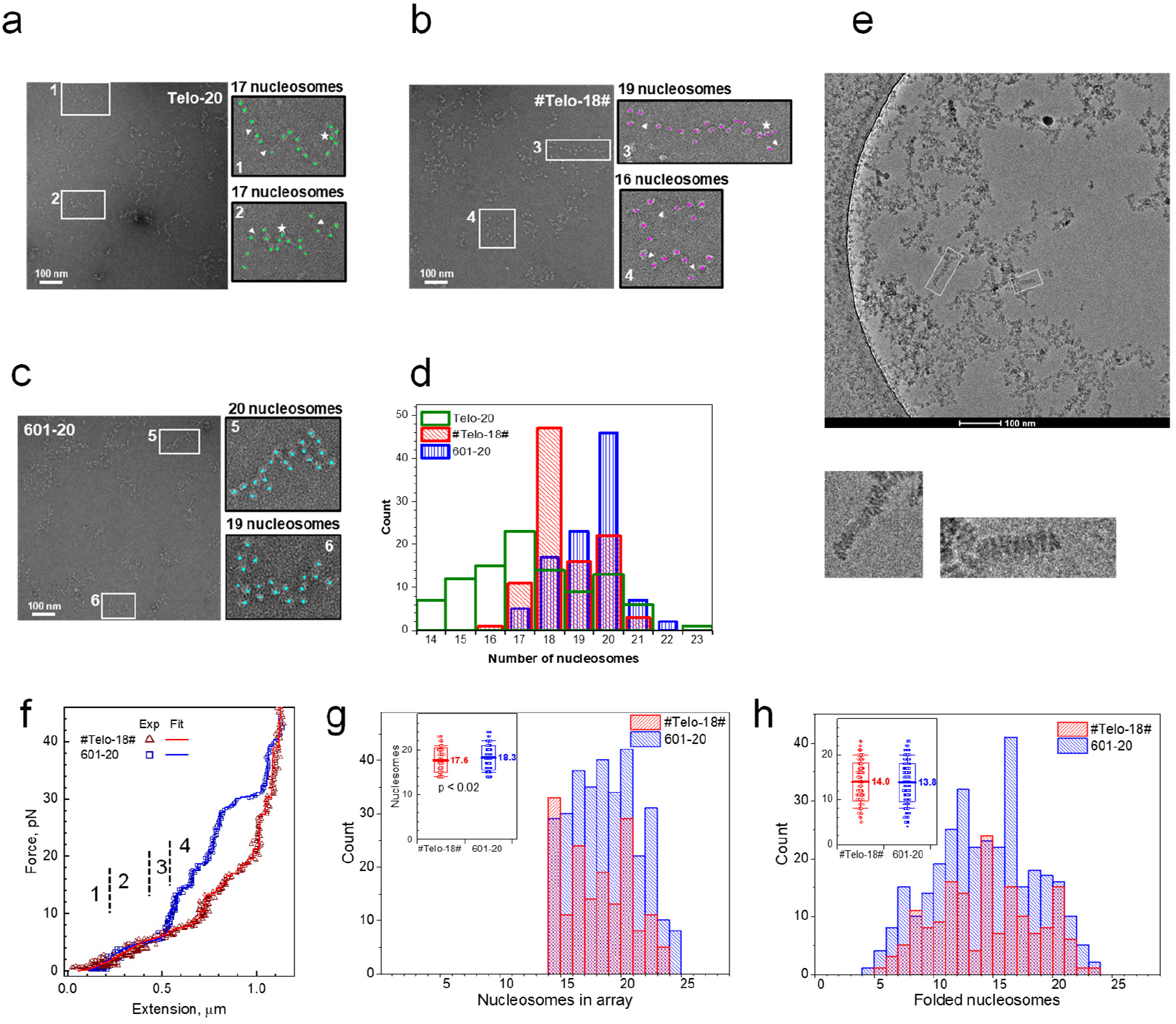
Heterogeneous nucleosome occupancy of telomeric chromatin. **a**, Representative negative stained EM micrographs at low salt conditions for **a** Telo-20, **b** #Telo-18# and **c** 601-20. Stars and triangles show respectively narrow and wide nucleosome spacing. **d,** Nucleosome count in arrays (arrays < 14 nucleosomes excluded). **e,** Cryo-EM images of reconstituted telomeric arrays were obtained from a mixture of ligated DNA templates (Extended Data Fig. 2f and k). **f,** Examples of force-extension curves for #Telo-18# (red) and 601-20 (blue). Numbers refer to respective transitions in the model describing the stretching. **g,** The number of nucleosomes in #Telo-18# (red) and 601-20 (blue) is determined by the MMT method. **h,** Number of folded nucleosomes in #Telo-18# (red) and 601-20 (blue). Inserts in **g** and **h** show respective statistics (traces < 14 nucleosomes excluded).

The telomeric arrays were also analysed by analytical ultracentrifugation sedimentation velocity (AUC-SV) analysis, which confirmed the heterogeneity of the telomeric arrays (Extended Data Fig. 3a). We conclude that under low salt conditions, *in vitro* reconstituted telomeric chromatin fibres are characterised by a variable nucleosome occupancy with irregular nucleosome spacing. These results confirm previous observations that nucleosomes formed on telomeric DNA are mobile and less stable than fibres formed on ‘601’ DNA templates^19, 22^.

## The compaction of telomeric chromatin display a columnar arrangement

Since telomeric DNA lacks nucleosome positioning information, we asked how this affects the compaction of telomeric chromatin fibres. We hypothesized that this unique feature might result in novel and different structural characteristics.

We visualized fibre folding of the 20-mer telomeric nucleosome arrays, which were compacted using standard conditions in the presence of Mg^2+^, with negative stain EM^15, 22^. Interestingly, among irregularly compacted globular fibres, we observed many arrays containing stretches of well-defined columnar structures of stacked nucleosomes that persist over at least four nucleosomes (Extended Data Fig. 3b, c). For the 601-20 array, ladder-like fibres dominated the well-defined particles (Extended Data Fig. 3d), in agreement with a previous study of the 601-157 nucleosome arrays. Notably, the columnar conformation was specific to the telomeric fibres and absent for the 601-20 arrays. A columnar form of telomeric chromatin was previously proposed^24^.

To further investigate the compaction of telomeric chromatin fibres and establish whether shorter arrays display more well-defined particles, we investigated the Mg^2+^ induced folding of nucleosome arrays formed on DNA templates generated by self-ligation of 4 repeats of 157 bp telomeric DNA (Extended Data Fig. 2f and k) using Cryo-EM. Remarkably, almost all arrays (98%) displayed distinct compact columnar structural features (Fig. 1e). Under identical folding conditions, the 4×157 bp Widom ‘601’ tetranucleosome compacted into a zig-zag structure that appeared similar to the one observed previously in the crystal structure of the 157-601 tetranucleosome (Extended Data Fig. 4a)^14^.

## Single-molecule force spectroscopy reveals the mechanical properties of telomeric chromatin fibres

Next, we characterised the mechanical and folding properties of the telomeric chromatin fibres by measuring the response of the fibres to single-molecule stretching with force spectroscopy. Multiplexed magnetic tweezers (MMT) were used to investigate the mechanical properties of the telomeric hybrid #Telo-18# in comparison with the 601-20 arrays under folding conditions in the presence of physiological Mg^2+^ concentration (see Methods).

Examples of force-extension curves recorded for the #Telo-18# and 601-20 arrays (Fig. 1f) display the characteristic transitions observed during force-induced nucleosome array unwinding (Extended Data Fig. 3e and f)^16^: (1) Extension of the folded array characterised by the stretching modulus, *k_fibre_*. (2) Transition the fibre to a beads-on-a-string chain accompanied by nucleosome unstacking and partial DNA unwinding characterised by a free energy *ΔG_1_*. (3) Deformation of the nucleosomes with further DNA unpeeling characterised by free energy *ΔG2*. (4) Irreversible one-step rupture of the last turn of the DNA. Data were collected and analysed, giving the total number of nucleosomal particles (nucleosomes, hexasomes, and tetrasomes) (Fig. 1g) and the number of fully folded nucleosomes (Fig. 1h) in the #Telo-18# and 601-20 arrays. The results agree with our EM (Fig. 1d) and AUC (Extended Data Fig. 3a) data, displaying variation in nucleosome occupancy and a smaller number of nucleosomes in the #Telo-18# relative to 601-20 arrays. Interestingly, we observed that the mean numbers of stacked folded nucleosomes are similar for the 601-20 and #Telo-18# fibres (Fig. 1f). This suggests that the #Telo-18# arrays form dense structures with close nucleosome-nucleosome contacts, as observed in the EM images of the telomeric fibres (Fig.1e and Extended Data Fig. 3b, c).

Analysis was made to obtain the parameters describing the fiber mechanical properties (Methods). The fibre stiffness, *k_fibre_*, displayed somewhat lower values for the telomeric chromatin (*k_fibre_* = 0.49 pN for #Telo-18# and *k_fibre_* = 0.53 pN for 601-20) (Extended Data Fig. 3g). Similarly, the free energy, *ΔG_1_*, is slightly lower for the telomeric fibre (12.6kT for #Telo-18# and 13.6kT for 601-20) (Extended Data Fig. 3h). The free energy *ΔG2* is similar for both arrays (Extended Data Fig. 3h). Compared to previously characterised fibres with longer NRLs, *ΔG_1_* is smaller, *ΔG2* is similar, and *k_fibre_* is between the values found for the 167 and 197 NRL 601 fibres under similar conditions^25^. Though not conclusive, such values would be consistent with an array of nucleosomes with very short linker DNA and weak stacking between nucleosomes. Analysis of the rupture corresponding to the final stretching of the fibre^26^ indicated weaker DNA binding to the histone octamer in the #Telo-18# nucleosomes relative to the 601-20 array (Extended Data Fig. 3i,j) in agreement with the weak HO affinity to telomeric DNA^4^.

## The telomeric tetranucleosome condenses into a column of tightly stacked nucleosomes

We focused our Cryo-EM structural analysis on telomeric tetra-nucleosomes folded in the presence of Mg^2+^. A Cryo-EM dataset of “Telo-tetra” was processed (Fig. 2a-c, Extended Data Table 1, and Extended Data Fig. 4b,c) to 8.1 Å resolution (Extended Data Table 1, Extended Data Fig. 2a, Extended Data Fig. 5, and Extended Data Fig. 7a and b). Remarkably, the Telo-tetra structure clearly exhibited four nucleosomes stacked on top of each other with DNA in a continuous one-start left-handed spiral (Fig. 2a-c). Four 145 bp telomeric mononucleosome X-ray structures^19^ were used to fit into the EM map (Fig. 2b). The rigid fit gave significant overlap for the DNA ends, which were trimmed and rebuilt to fit into the EM density (Fig. 2c and Extended Data Table 2). No distinct region can be demarcated as linker DNA with adjacent histone octamers concomitantly binding DNA at the interface of the two octamers. Furthermore, the tetranucleosome columnar packing was confirmed in three different and independent datasets. Analysis of a segmented tri-nucleosome (Tri-NCP) unit strongly suggested that the stacked di-nucleosome is the asymmetric unit in the columnar structure observed (Extended Data Table 1 and 2, Extended Data Fig. 4d).

**Fig. 2.**
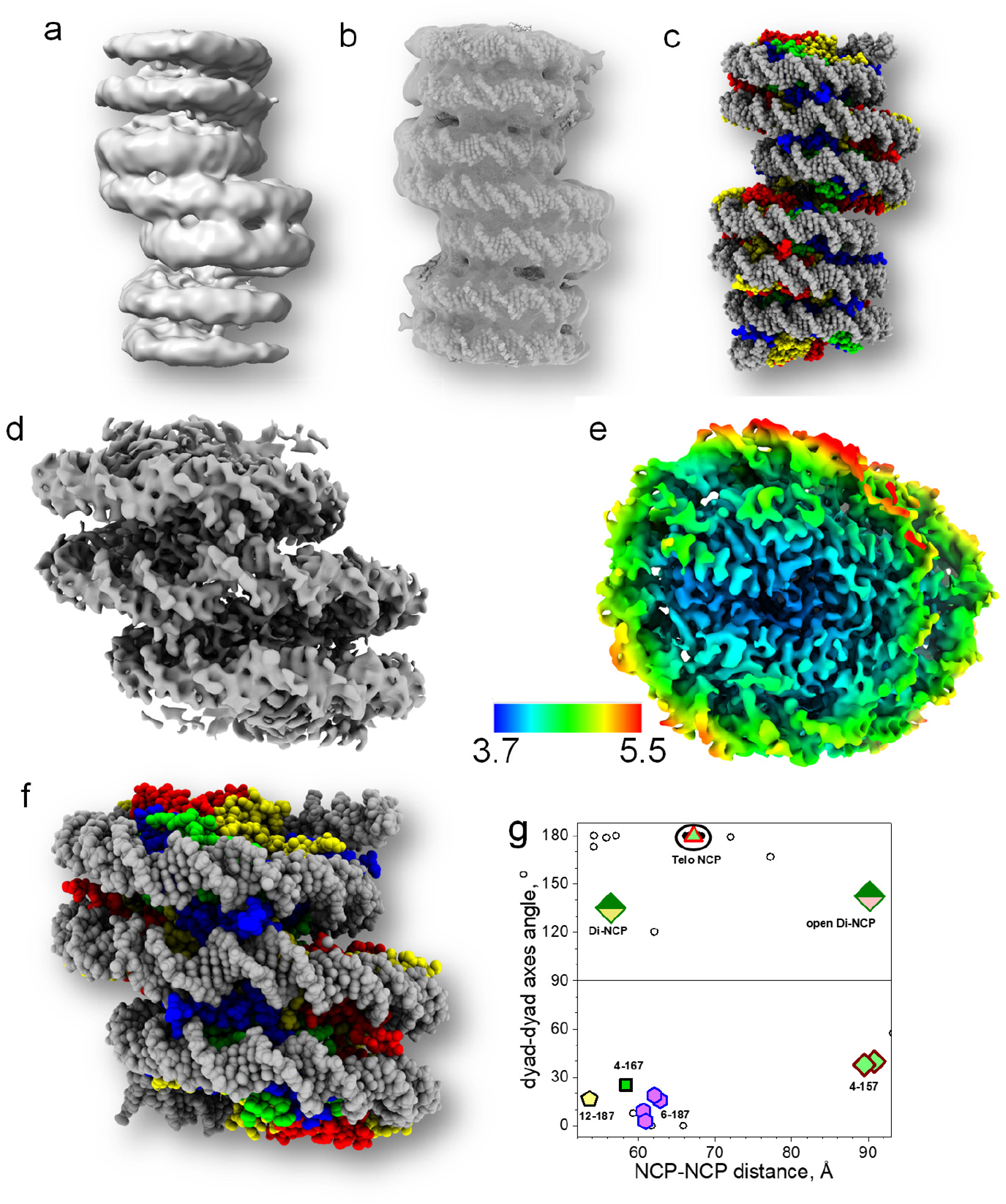
Structure of the telomeric tetranucleosome. (In all panels described in Fig. 2-4, when coloured, H3, H4, H2A, and H2B are shown as blue, green, yellow, and red, respectively. DNA is coloured grey.) **a,** EM map of the Telo-tetra. **b,** Four telomeric NCPs fitted into the map of the Telo-tetra array. DNA is displayed as spheres and protein residues as balls and sticks. **c,** The telomeric tetranucleosome structure. The DNA and histones are shown as space-filling balls. **d,** Electron density map and **(e)** local resolution in the Di-NCP map. **f,** The di-NCP structure was obtained from fitting into the map. **g,** Correlation between NCP-NCP distance and dyad-dyad angle. The telomeric di-nucleosome structures are shown by green-yellow (Di-NCP) and green-pink (open Di-NCP) rhombi. Small hollow circles represent data in NCP crystals. The red-light green triangle is from the recent Telo-NCP crystal structure^19^. Data from published nucleosome arrays: 4-157^14^ (blue-green rhombi), 4-167-601^41^ (blue-green square), 6-187-601^12^ (blue-magenta hexagon) and 12-187-601^13^ (black-yellow pentagon).

We next processed particles using a box size characteristic of the di-nucleosome (Di-NCP) sub-structural unit in a separate dataset (Extended Data Table 1), which gave a Di-NCP map at 3.9 Å resolution with well-resolved helices and DNA base separation at the interface of two NCPs (Fig. 2d and e, Extended Data Fig. 4e, Extended Data Fig. 7g and h; Extended Data Table 2). Two 145 bp telomeric NCP X-ray structures^19^ were fitted into the EM density, followed by rebuilding histone tails and the DNA (Fig. 2f and Extended Data Table 2).

We compared the stacking distances and NCP-NCP orientation in the NCPs of the columnar model with chromatin fibres from X-ray and EM models (Fig. 2g)^27^. In the telomeric Di-NCP, the parameters characterising the nucleosome stacking geometry showed significant deviations compared to the stacking observed in most published X-ray and Cryo-EM structures of nucleosome arrays^12–14, 28^. Furthermore, the short NRL results in a near head-to-tail arrangement of NCPs as defined by the dyad axes of the stacked telomeric NCPs (about 130°), in contrast to most other chromatin fibres characterised by a head-to-head orientation (close to 0°) (Fig. 2g). The short NCP-NCP distance of 56 Å in the telomeric Di-NCP contrasts with that observed between the N and N+2 nucleosomes in the two-start 157-601 tetranucleosome 90 Å^14^.

## Histone tails mediate and stabilize the columnar stacking

To establish the role of the histone tails in NCP stacking, we fitted the telomeric X-ray NCP structure into a Di-NCP map refined from three merged datasets and built tail densities (Fig. 3, Extended Data Table 1, Extended Data Table 2, Extended Data Fig. 5, and Extended Data Fig. 6, Methods). The identified tail densities are well defined in the three independent datasets that we processed. Given the present resolution and the expected dynamic feature of the tail interactions, only a minority of tail side chains can be assigned. However, a significant fraction of the tail densities can be confidently identified by starting at the root of the tail following the path of the tube-like densities in the map (Extended Data Fig. 8a).

**Fig. 3.**
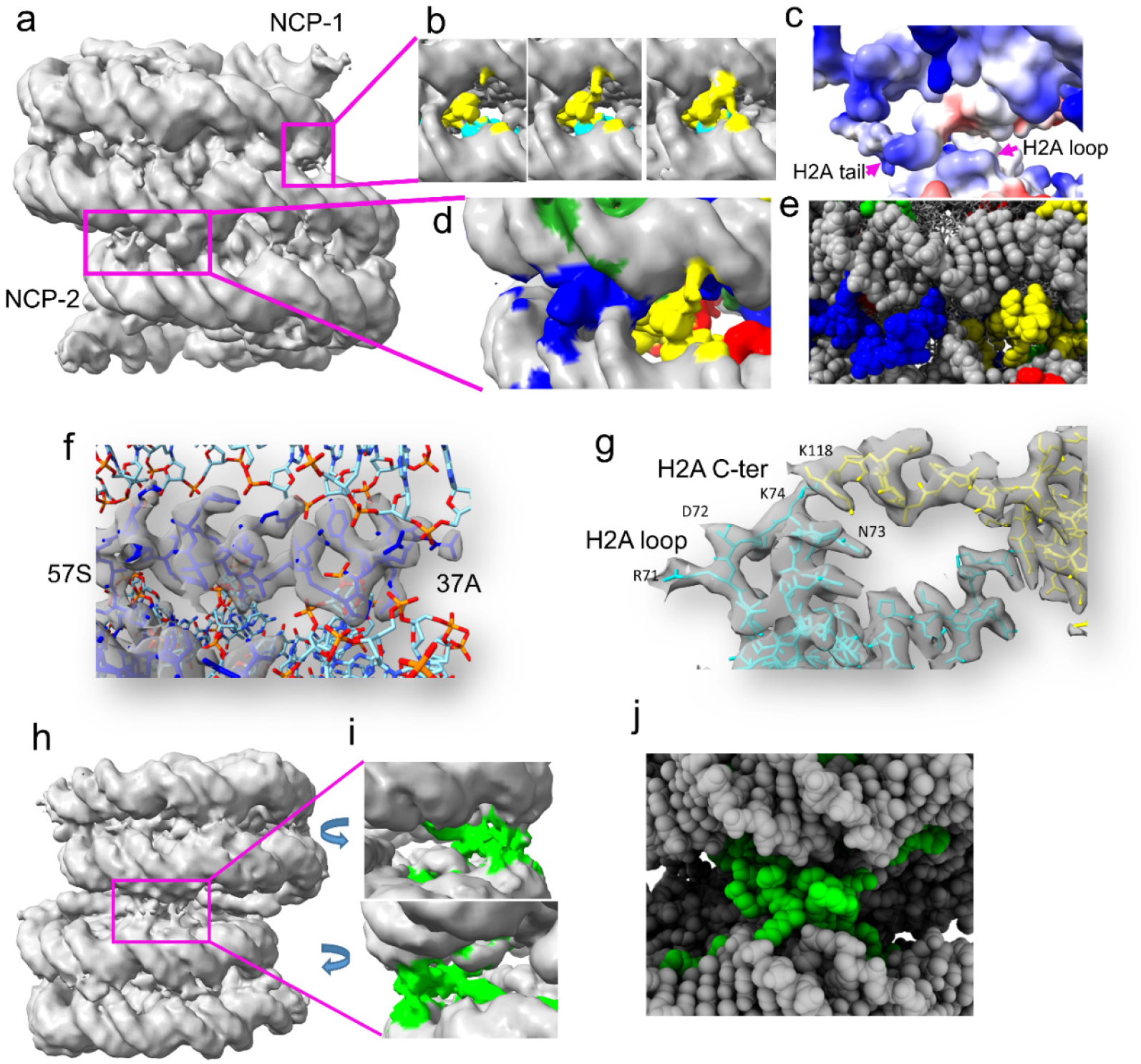
Histone-mediated interactions stabilizing the columnar tetranucleosome/Di-NCP structure. **a,** Clamp formed by the H2A C-terminal tails and the H3 N-terminal helix. **b,** Close-up view of the EM density corresponding to the H2A mediated NCP-1-NCP-2 interaction in (a) (small magenta box). Panels show maps with decreasing contour density from left to right. **c,** H2A C-terminal interaction with the H2A loop is suggested to be mediated by a patch of polar residues that forms a weak acidic patch-basic patch interaction. **d,** Density corresponding to the H3 N-terminal helix-H2B C-terminal clamp interacting with DNA. **e,** H3-H2A clamp as a space-filling model. **f,** Amino acid residues 37-57 of the H3 N-terminal helix (blue). **g,** Amino acid residues of the H2A loop (cyan) and H2A C-terminal (yellow). **h,** Map depicting the location of H4 tails stabilizing NCP-NCP stacking. **i,** Density of the H4 tails of chains of B and P. **k,** The H4 tails built into the EM density are shown as balls (green).

The N-terminal helices of H3 show strong density along the minor groove at the boundary of NCP-1 and NCP-2 and act as anchor points for the DNA (Fig. 3a, d-f). The C-terminal tail of H2A mediates the second set of interactions (Fig. 3a-d and g). There is a continuous density along loop 2 of H2A from the adjacent NCP (Fig. 3b, c, d, and g) bridging the two NCPs. At lower contour, the map shows a continuous density that originates primarily from H2A K118, with minor contributions from the adjacent patch of polar residues S122, E121, T120 of the C-terminal tail congruous with R71, D72, N73, K74, K75 of the H2A loop 2. These interactions are complemented by the H2A C-terminal tail interaction with its DNA and with the DNA of the adjacent NCP, forming a multivalent bridging tether that facilitates the NCP-NCP stacking (Fig. 3b and d). We tentatively assigned an additional set of bridging interactions mediated by H4 tails (Fig. 3h-j). We noted that the H4 tail position also suggests a putative electrostatic interaction with the H3 core (Extended Data Fig. 8e).

The two H3 N-terminal helices and the H2A C-terminal tails occupy a crucial position in the region where the transition occurs between NCP-1 and NCP-2. Two regions are separated by 180° on the Di-NCP structure, where three DNA gyres are in the same plane (Fig. 3a and h). The H3 N-terminal helix and the stretch of positive residues (aa 124-129) of the H2A C-terminal tail appear to form a DNA clamp along the two minor grooves through interactions with the DNA phosphates (Fig. 3d and e). We suggest that all these tail domains form a network of synergistic interactions that facilitate and mediate the stabilization of the columnar stacking.

## The columnar form of telomeric chromatin displays an unusually short nucleosome repeat length

We calculated the dyad-to-dyad distance to estimate the NRL in the telomeric structures. The NRL is ∼132±2 bp for all the Di-, Tri- and Tetra-nucleosome structures. Digestion with MNase resulted in bands corresponding to ∼130 and ∼157 bp (Extended Data Fig. 4f). The ∼132 bp NRL calculated from the columnar structure is consistent with the in vitro and in vivo nuclease assays (Extended Data Fig. 4f), suggesting that our *in vitro* structure is consistent with the *in vivo* assays. Furthermore, our structure implies that the 157 bp band arises from the 132 bp DNA protected by the 14 distinct histone-DNA contacts and the adjacent DNA protected by the H2A C-terminal tails, which are ∼157 bp apart in the columnar structure (Extended Data Fig. 9d, e). We suggest the ∼130 bp band results from elevated nucleosome breathing on the telomeric NCP and the observed spontaneous unwrapping of DNA ends in 10 bp segments^29, 30^. Additionally, an ∼132 bp repeat length from *in-vivo* data for native chromatin has been previously observed, and a similar columnar packing was proposed^31^.

We then asked what is the DNA wrapping length on the HO in the sub-class of the Telo-tetra particles that can be identified as mononucleosomes. To investigate this, we picked and processed segmented isolated mononucleosome units in Telo-tetra particles characterised by an open state devoid of nucleosome-nucleosome interaction, resulting in a resolution of 3.5 Å and DNA length of 145±2 bp (Fig. 4 a-c and Extended Data Fig. 7i and j). This confirms that the short NRL of 132 bp is unique to the columnar tightly stacked telomeric fibre.

During 3D classification, we also noted a well-defined class that showed two NCPs near 90° orientation relative to each other, which was also observed as distinct 3D classes among tri-NCP and Telo-tetra particles (Extended Data Fig. 4d and Extended Data Fig. 9a-c). We designate this di-nucleosome as ‘open Di-NCP’ (Extended Data Fig. 6). We processed the volume to a resolution of 4.5 Å (Fig. 4d-f, Extended Data Fig. 9a, and Extended Data Fig. 7 k and l). Comparison of the stacked and open Di-NCP structures showed that they have similar head-to-tail dyad axes orientations (Fig. 2g), but the plane-plane angle between the NCP flat cylinders in the open Di-NCP is close to 90° compared to 22° for the stacked conformation. We suggest that this structure corresponds to a stable intermediate state in the folding (or unfolding) of Telo-tetra (Extended Data Fig. 9b and c). A recent Cryo-EM structure of the archaesome observed a similar feature^32^. The lower resolution (Extended Data Fig. 9a) for one of the NCPs in the open Di-NCP indicates an average structure with multiple HO conformations and subsequent divergent DNA phasing.

**Fig. 4.**
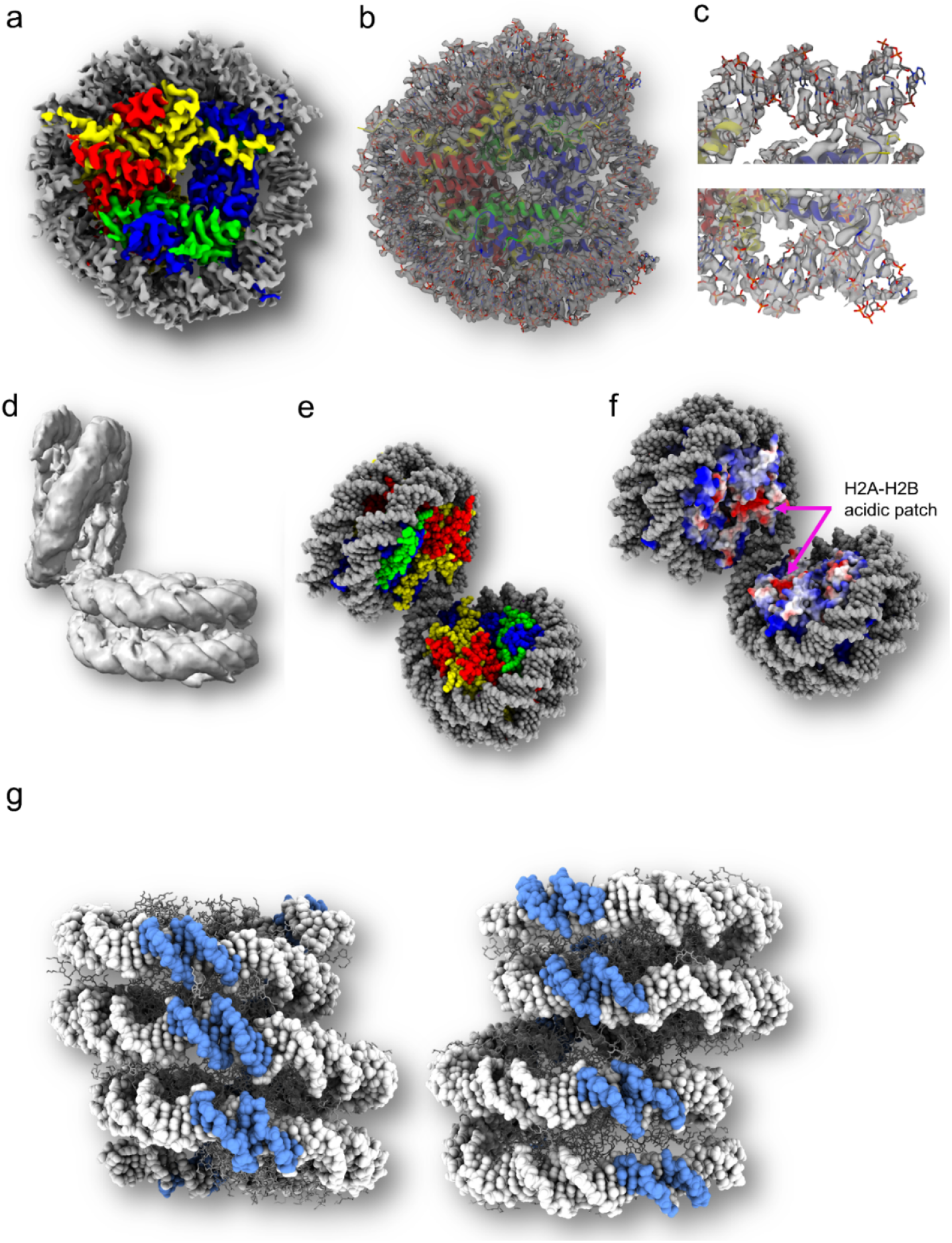
Features of telomeric chromatin. **a,** Mononucleosome structure in unfolded Telo-tetra particles. **b,** A 145 bp telomeric mono-NCP fitted into the map. The histones are shown as a cartoon. **c,** Fitting of the DNA ends of the 145 bp telomeric NCP. **d,** Map of the open Di-NCP. **(e)** Structure of the open Di-NCP rebuilt from two mono NCP. **f,** Exposed H2A-H2B acid patch with the histone core coloured corresponding to surface charge. **g,** An illustration of super-grooves spanning multiple nucleosomes in Telo-tetra.

The unique columnar arrangement of the telomeric tetranucleosome also results in the alignment of multiple DNA minor grooves, creating a continuous ‘super-groove’ ^33^ along the nucleosome column (Fig. 4g), resulting in the alignment of two pairs of three minor grooves across two nucleosomes (Fig. 3a and h).

## Conclusions

This work revealed the columnar organization of telomeric chromatin, and its features suggest significant structural insights of importance for telomere function *in vivo*. It has been shown *in vivo* that DDR at the telomere does not require decompaction^34^. SIRT6, which plays a multifaceted role in maintaining telomeres through its deacetylase activity, activation of DDR factors, recruitment of remodeller and helicase^35^, like many other factors, engages the H2A-H2B acidic patch for nucleosome binding^36, 37^. Importantly, the intermediate open Di-NCP structure exposes the buried acidic patch (Fig. 4f, Extended Data Fig. 8b-d) and provides a structural basis for binding such factors to the acidic patch within the context of the telomere without the need for telomeric heterochromatin decompaction. The presence of the open structure in the archaeal^32, 38^ chromatin also suggests an evolutionarily conserved function served by this stable intermediate in the organization and maintenance of chromatin^32^.

Our structure highlights the pivotal role of the H3 N-terminal helix and H2A C-terminal tail in mediating its dense stacking. This sheds light on the effect of recruitment of H2AX during DDR at the telomere^5–8^. H2AX lacks K125 and K129 at the C-terminal tail, which contacts the DNA in our structure. Thus, incorporating H2AX that is lacking these positive residues will destabilize the structure and facilitate the recruitment of protein factors, including DDR proteins.

We speculate that the super grooves identified on Telo-tetra serve two functions. First, the supergroove spanning multiple nucleosomes is ideal for TRF1 and TRF2 that binds as homodimers, facilitating capping and compaction of telomeric chromatin^39^. Secondly, the supergroove may facilitate chromatin remodellers that engage multiple DNA gyres on nucleosomes for binding^40^. Finally, in the columnar structure, DNA is exposed on the surface, and this may contribute to the susceptibility of telomeric DNA to damage.

The knowledge of the organization of telomeric chromatin at the molecular level revealed in this work paves the way for structural studies involving telomeric binding factors, the interaction with shelterin components and DDR proteins. Such information will help us shed light on the structure-function relationship of telomeres in the context of ageing and cancer. Our columnar fold resembles that formed by archaeal histones and a partial dinucleosome structure with a nucleosome repeat length ranging from 125-147 bp^32, 38^. Thus, it is plausible that a columnar structure is present in chromatin regions outside the telomere as previously proposed^31^.

## Methods

### Construction of pTelo-1 vector

A 151 bp human telomeric ssDNA fragment (C-rich strand) flanked by two custom-designed multiply cloning sites, MCSs (Extended Data Fig. 1a), were purchased as a PAGE Ultramer^®^ DNA Oligo (IDT, USA). The synthetic ssDNA oligo was converted to dsDNA by primer extension reaction at 60° C using iProof DNA polymerase (Bio-Rad, USA). The resulting duplex was cloned between the EcoRI and HindIII sites of the pUC57 vector. The ligation product was transformed into *E. coli* SURE 2 Supercompetent cells (Stratagene, USA) for clonal selection and amplification. The construction of the resulting plasmid, termed pTelo-1 (Extended Data Fig. 1a), was confirmed by restriction profile analysis and sequencing of the telomeric insert. To limit undesired DNA rearrangement events, pTelo-1 and its derivative constructs were amplified in SURE 2 *E. coli* strain and cultured at 30°C in LB medium at low cell density (OD_600_ < 2). Milligram quantities of the various plasmids used in this work were purified using the Plasmid Giga Kit (QIAGEN, Germany) according to the manufacturer’s instructions.

### Construction of short telomeric array templates

In pTelo-1, the two custom-built MCSs flanking the telomeric DNA host various endonuclease restriction sites, enabling various manipulations, modifications, and release of the telomeric insert (Extended Data Fig. 1a). Two AvaI sites are contiguous to the telomeric insert (one on each side). Digestion of pTelo-1 with AvaI releases single telomeric units that can be subsequently multimerized into short DNA arrays by ligation (Extended Data Fig. 1b). The resulting arrays consist of integral multiples of 157 bp telomeric units. The non-palindromic property of the AvaI overhangs ensures control over strand orientation. Briefly, a purified AvaI-released Telo-1 DNA fragment was self-ligated overnight at 4°C. The polymerized mixture was re-ligated into the parental backbone, and the reaction products were transformed into *E. coli* SURE 2 competent bacteria for clonal selection and amplification. The pTelo-4 construct, containing four 157 bp telomeric DNA units, was identified following screening by partial AvaI digestion and sequencing of the telomeric array. Hereafter, all the plasmids carrying telomeric array templates are designated with the pTelo-*N* symbol followed by an integer (*N*) denoting the number of 157 bp units in the array.

### Construction of intermediate telomeric array templates

Construction of longer telomeric DNA arrays relies on two Type IIS restriction enzymes BbsI and BtgZI, which cut the telomeric constructs within the utmost upstream and downstream AvaI sites, respectively (Extended Data Fig. 1a). Unlike AvaI, the former endonucleases let unilateral modifications of the construct or release entire telomeric arrays bearing AvaI-compatible overhangs. Such an approach was applied to construct pTelo-8 from pTelo-4 (Extended Data Fig. 1c). Briefly, the parental construct containing a telomeric array was linearized with BbsI and dephosphorylated using rSAP. Concurrently, a telomeric array was excised from the parental vector by EcoRV digestion, dephosphorylated, and fractionated by agarose gel electrophoresis. The resulting purified telomeric insert was further digested with BtgZI to produce a distal AvaI overhang, ligated to the BbsI-linearized parental construct, and purified by agarose gel electrophoresis. Finally, the resulting purified product was deprotected with BbsI yielding a proximal AvaI overhang and enabling the circularization of the resulting plasmid. As for the previous telomeric constructs, the integrity of the telomeric arrays obtained by this method was verified by partial AvaI digestion and restriction profile analysis. However, sequencing the complete array was impossible due to the repetitive nature of the telomeric sequence and its length (>1 kb). Similarly, constructs hosting telomeric arrays ranging from eight (pTelo-8) to twelve (pTelo-12) units were successfully produced using the same method. However, longer telomeric arrays proved to be very unstable during their amplification in *E. coli*. The method was also applied to construct an array of twenty repeats of a 157 bp Widom 601 nucleosome positioning DNA (601-20, Extended Data Fig. 2a and S2e).

### Construction of long telomeric array templates

Long telomeric DNA arrays were obtained by large-scale ligation of two intermediate arrays (Extended Data Fig. 1d). Succinctly, 10-100 mg pTelo-10 vector was digested with EcoRV to release its 10×157 bp telomeric DNA array. The EcoRV reaction mix was supplemented with alkaline phosphatase and DdeI, DpnI, and HaeIII endonucleases to fragment the 2647 bp pTelo-10 backbone into small (< 370 bp) pieces that can be readily removed by fractional PEG precipitation. Subsequently, the Telo-10 DNA fragment was further purified by size exclusion chromatography on a Sephacryl S-500 column and eluted in 50 mM Tris-HCl (pH 8.0), 5 mM Na-EDTA and 0.5 M NaCl. The resulting purified Telo-10 array was separately digested with either BtgZI or BbsI to generate AvaI overhangs at the downstream and upstream extremities, respectively, followed by purification by 8% PEG precipitation. Finally, BtgZI- and BbsI-digested fragments were combined at an equimolar ratio and ligated (4°C, 16 h). The resulting Telo-20 ligation product was further purified by phenol-chloroform extraction. Typically, the applied experimental conditions yielded about 75% ligation efficiency (Extended Data Fig. 2c). This method proved suitable for preparing long array templates that would otherwise be too unstable when propagated in *E. coli*.

### Construction of long hybrid telomeric/601 array templates

A hybrid telomeric/601 DNA array template (#Telo-18#, Extended Data Fig. 2a and d) consisting of 18×157 bp telomeric DNA units flanked by a 157 bp Widom 601 fragment (denoted as #) at each terminus was prepared by large-scale ligation of #Telo-8 and Telo-10# DNA arrays. #Telo-8 and Telo-10# DNA arrays were respectively built by extending the upstream side of a Telo-8 array and the downstream side of a Telo-10 array with a 1×157 bp Widom 601 insert. The resulting constructs (termed p#Telo-8 and pTelo-10#, respectively) were used to release the aforementioned #Telo-8 and Telo-10# DNA arrays.

### Large-scale amplification and purification of Telo-4 DNA array templates

The telomeric tetranucleosome DNA template (Telo-4 from pTelo-4) and 601 tetranucleosome template was amplified and purified as previously described^19^. Briefly, pTelo-4-transformed *E. coli* SURE 2 were grown in a TB medium at 30°C. The plasmid was extracted by standard alkaline lysis^42^, and the Telo-4 array template was released by EcoRV. The vector backbone was partially removed by Mg^2+^-mediated PEG size fractionation, followed by ion-exchange chromatography. The Telo-4 DNA was eluted using a 300-750 mM LiCl gradient buffer (20 mM Tris-HCl (pH 7.5) 1 mM Li-EDTA). The multimers of Telo-4 were prepared by large-scale self-ligation of EcoRV-released Telo-4 DNA insert (Extended Data Fig. 2f).

### Human histone octamer preparation

Recombinant human histone proteins were individually expressed, purified, and refolded into human histone octamers essentially as previously described^43, 44^. Briefly, individual histone proteins (H2A, H2B, H3, and H4) were mixed under high salt at a ratio of 1.2:1.2:1.0:1.0, and the assembled octamer was purified by size exclusion chromatography.

### Telomeric nucleosome array preparation

The 20-mer DNA and recombinant human histone octamers were reconstituted into nucleosome arrays by the salt dialysis method, essentially as described previously^23^. Salt concentration was reduced from 2 M to 2.5 mM LiCl by continuous dialysis at a 0.6 mL/min flow rate overnight using a peristaltic pump. The final buffer consisted of 10 mM HEPES-LiOH pH 7.4, 1 mM Li-EDTA, 1 mM DTT, and 2.5 mM LiCl. The saturation level of the reconstituted arrays was assessed by 0.2×TB-agarose gel electrophoresis, stained with SYBR™ Gold Nucleic Acid Gel Stain (ThermoFisher Scientific, USA). All Cryo-EM samples were reconstituted employing continuous dialysis from 2 M LiCl to ∼65 mM LiCl in buffer containing 20 mM Tris-HCl (pH 7.5), 1 mM EDTA and 1 mM DTT across 16 hrs. The sample was further dialyzed in 20 mM Tris-HCl, 1 mM EDTA, and 1 mM DTT for 4 hrs before centrifugation and harvest.

### Negative stain Electron microscopy

All samples visualized with negative stain EM were fixed with 0.02% (v/v) glutaraldehyde on ice for 10 minutes, at a concentration of 30-40 µg/mL. 4 µl of the sample was applied on a carbon-coated grid for 1 minute and subsequently blotted from the edge of the grid using a Whatman Grade 1 filter paper. Immediately, 4 µL of 2% (w/v) uranyl acetate was applied on the grid and left to stain for another minute. The grids were visualized with a Tecnai T12 electron microscope operating at 120 kV, and micrographs were recorded with an Eagle 4 K CCD camera (FEI) at 49,000× magnification with a defocus value of −1.2 μm.

Extended nucleosome arrays were diluted to the desired concentration using the final reconstitution buffer (10 mM HEPES-LiOH pH 7.4, 1 mM Li-EDTA, 1 mM DTT, and 2.5 mM LiCl). From each sample, 100 arrays that displayed a bead-on-the-string conformation had their nucleosome occupancy quantified, based on a threshold of 14 nucleosomes per array. This threshold allowed to omit arrays from unligated DNA fragments during the counting process.

The compacted arrays were prepared by overnight dialysis of the nucleosome array against the reconstitution buffer supplemented with either 0.6 or 0.8 mM MgCl_2_. Both telomeric arrays compact and precipitate at a lower Mg^2+^ concentration, with an EC_50_ value of 1.05 mM ± 0.02 mM. Compaction of telomeric arrays was achieved with the addition of 0.6 mM MgCl_2_ during the dialysis. In contrast, 601-20 array precipitates at higher Mg^2+^ concentration (EC_50_ = 1.34 mM ± 0.02 mM) and 0.8 mM MgCl_2_ was used to induced array compaction. The concentration of Mg^2+^ was determined from the Mg-precipitation assay, at which 85% of the arrays remain in the soluble fraction. EC_50_ refers to the average concentration of Mg^2+^ at which 50% of the array was precipitated.

### Cryo-Electron microscopy

Most published literature probing for ordered chromatin structures *in vivo* employs fixation by glutaraldehyde^45^ to preserve the structure. In our experience with telomeric nucleosomes, the telomeric nucleosome dissociated at the glutaraldehyde concentration used in such studies. Also, it has been shown that glutaraldehyde perturbs nucleosome and bias structure^46^. We screened various buffers and ionic conditions to obtain uniform homogenous samples to preserve structural features without fixing the sample. Potassium cacodylate buffer at pH 6 supplemented with 0.02-0.2 mM Mg^2+^ gave the best-preserved Telo-tetra on the EM grid. The low pH decreases the DNA’s solubility and adds ∼18 positive charges to the histone octamer. The higher positive net charge on the histone octamer facilitates a tighter binding of the DNA strand, thereby decreasing the DNA dynamics, strengthening the histone-DNA interaction, and stabilizing the overall structure of the condensed tetra nucleosome. The unusually low Mg^2+^ concentration required to compact the array is due to the short 10 bp DNA linker and the sub physiological pH of 6.0. To eliminate any artefacts that may arise from the use of cacodylate buffer at pH 6.0, we screened a 157 bp 601 tetra nucleosome under identical conditions and obtained arrays showing the conventional zig-zag arrangement in agreement with the literature (Extended Data Fig. 7a)^14^.

Tetranucleosome arrays in 20 mM Tris pH 7.5, 1 mM Li-EDTA, and 1 mM DTT were buffer-exchanged into cacodylate buffer (pH 6.0) containing 20 mM potassium cacodylate and 1 mM DTT using Amicon centrifugal filter (cutoff 10 kDa). The array sample was subsequently diluted (1:1) with cacodylate buffer containing appropriate amounts of Mg^2+^ to give a sample concentration of 1 mg/ml arrays in 20 mM potassium cacodylate, 0.04-0.2 mM Mg^2+^ and 1 mM DTT. This sample (4 µL) was applied to a freshly glow discharged quantifoil grid (R1.2/1.3 Cu 200 mesh) at 4°C at 100 % humidity, blotted (time: 2 s, blot force: 1, drain: 1 s), and plunge frozen (FEI vitrobot) in liquid ethane.

Data collection was carried on two Titan KRIOS microscopes (Thermo Fischer Scientific) equipped with K2 and K3 direct electron detectors in counted mode. Three datasets are presented here, of which two (Dataset 1 and 2) were collected at 1.4 Å/px and the third one (Dataset 3) at 0.86 Å/px. Micrographs were collected at a magnification of 105,000× corresponding to a pixel size of 1.4 Å (K2)/0.86 Å (K3) in movie mode with a total dose of 50 e/Å^2^ at defocus values −0.5, −1.0, −1.5, −2.0, and −2.5 µm. The data processing was carried out using RELION and XMIPP in the SCIPION package^47^. Selected micrographs were retained after motion correction with RELION implementation of motioncor^48^ and CTF estimation with CTFFIND4^49^. For Dataset 1, a small set of particles (box size-220-240 Å) were picked employing XMIPP3 manual picking, followed by XMIPP3^50^ automated picking to pick all coordinates. Multiple rounds of 2D classification were carried out to select coordinates corresponding to particles with two or more nucleosome stacks. The particles were re-extracted with a box size of 240 Å, 314 Å, and 336 Å corresponding to Di-NCP, Tri-NCP, and Telo-tetra, respectively. The initial model for the template for 3D classification was made with a RELION initial model with a small set of manually picked particles. The 3D classification was carried out on the selected set of particles, and selected volumes were refined, followed by Bayesian polishing and local resolution estimation in RELION. The maps were visualized in Chimera^51^ or ChimeraX^52^.

For Dataset 2 and Dataset 3, the picking was carried out employing a 3D model from Dataset 1 as a template. All three datasets confirmed the columnar packaging of Telo-tetra. However, the population of various compacted states showed variation between the three datasets. A higher Mg^2+^ (>0.08 mM) gave homogenous and highly compact columnar particles; however, the aggregation on the grid resulted in significant difficulties in the data processing. Hence, we had collected the three datasets at Mg^2+^ concentration (0.06-0.08 mM) that resulted in compact individual particles with minimal aggregation. A Di-NCP map was generated for each of the datasets. As individual datasets gave a limited number of particles of the various higher order structures, we merged and processed selected micrographs from Datasets 1 and 2 to obtain the maps for Telo-tetra and higher order open states of Telo-tetra (Extended Data Table 1).

The models were generated by the rigid fitting of 145 bp telomeric NCP^19^ in Chimera. All fitting of mono NCP showed a significant overlap of DNA ends. These overlapping DNA ends were trimmed and joined in COOT^53^ to form continuous Di-NCP, Tri-NCP, or Telo-Tetra DNA. The histone octamer tails were trimmed to the core, followed by rebuilding in COOT in CCPEM^54^. The three Di-NCP maps from three datasets were combined in PHENIX with a resolution cutoff of 4.5Å. This map was used to rebuild the histone and was refined in COOT employing the ‘Chain refine’ option. The side chains that were resolved in the 3.9Å Di-NCP map were further refined employing the 3.9A Di-NCP map. The H2A N (up to aa 10) and C-terminal tail generally showed extra density than the telomeric mononucleosome crystal structure. The H4 tails that mediated stacking showed tube-like densities (up to aa 16), whereas the H4 tails that were not involved in stacking interaction did not show discernable EM density beyond aa 20. The H3 and H2A N-terminal tails showed densities indicative of their interaction with DNA in multiple conformations (between DNA gyres and groove). However, lack of resolution, multiple conformations, and the subsequent inability to fit side chains did not permit the rebuilding of respective tails in the final model. A representative model following tube-like density is shown in Extended Data Fig. 8a. The structures were refined using Refmac5^55^ in CCPEM^54^, followed by geometry minimization, COOT^53^ refinement, and validation using PHENIX^56^.

### Analytical ultracentrifugation

Sedimentation velocity experiments were carried out using a Beckman XL-I analytical ultracentrifuge (Beckman Coulter, Fullerton, CA) with an 8-hole An-50 Ti analytical rotor. Nucleosome arrays in 10 mM HEPES-LiOH pH 7.4, 0.1 mM Li-EDTA, 1 mM DTT, and 2.5 mM LiCl were diluted to OD259= 0.6-0.8 and loaded in 12 mm double-sector cells fitted with quartz windows. The samples were spun at 12,000 rpm, and 40 scans were collected at 10 min intervals. The data was fitted to a continuous c(s) distribution model^57^ using SEDFIT software and normalized with GUSSI software^58^. The sedimentation coefficients were corrected to s20,w using a partial specific volume of 0.66 mL/g, calculated from the mass ratio of DNA and histone octamer, while density and viscosity were calculated from the buffer composition using SEDNTERP^59^. For each condition, 3 separate samples were prepared in triplicate.

### Multiplex Magnetic Tweezers (MMT) measurements

The homemade MMT setup and flow cells used in this work were assembled using the design developed in the laboratory of Chromatin Dynamics of Leiden Institute of Physics, Huygens-Kamerlingh Onnes Laboratory, Leiden University^60, 61^. The solution of anti-digoxigenin (Merck Millipore, USA; 300 µL at 1 µg/µL) was introduced into the flow cell and incubated for 2 hours at room temperature, followed by flushing 1 mL of passivation solution (3.6 % Bovine Serum Albumin (BSA) heat shock fraction, pH 7, ≥ 98% (Merck), 0.1% Tween 20 (Merck)) with subsequent storage at 4°C.

Two types of nucleosome arrays with twenty repeats of the 157 bp NRL were studied using the MMT method: 601-20 and #Telo-18#. The former was prepared by the stepwise array extension method, while the latter was used by the in vitro ligation method described in the previous section, with one modification. Instead of EcoRV, EcoRI+BtgZI and HindIII+BbsI were used for excising the arrays from their plasmids. This resulted in 5’-EcoRI and 3’-HindIII overhangs that were ligated to asymmetric short linkers (∼25 bp) to achieve torsionally free DNA constructs for the MMT measurement. The 5’ and 3’ linkers were terminally labelled with digoxigenin-11-dUTP (Roche) or biotin-16-dUTP (Roche), respectively.

Nucleosome array fibres were reconstituted using the salt dialysis method, and the stoichiometric amount of histone octamers was empirically determined for each reconstitution. Each type of the array, #Telo-18# and 601-20 was treated identically. Two preparations of the arrays were made, testing three different HO:DNA ratios in each of the reconstitutions: HO:DNA ratios were 0.95, 1.00, and 1.05 in one; and 0.9, 1.0, and 1.1 in the other reconstitution. The quality of reconstitution was checked on 0.7% agarose EMSA using freshly reconstituted nucleosome arrays.

The flow cell was washed from passivation solution with 1 mL of measurement buffer, MB (100 mM KCl, 2 mM MgCl_2_, 10 mM NaN_3_, 10 mM HEPES pH 7.5, 0.1% Tween 20, 0.2% BSA). To attach the biotin-labelled end of DNA to streptavidin-coated beads, 2 µL of vortexed 10 mg/mL 2.8 µm paramagnetic beads (Dynabeads M-280 Streptavidin, Thermo Fisher Scientific Baltics UAB, Norway) was added to 0.1-1.0 pg/µL solution of nucleosome arrays in 500 µL of MB followed by incubation for 10 min with gentle mixing. Then, 500 µL of MB with the beads and the arrays attached to them was pumped into the flow cell by a syringe pump (New Era Pump Systems, USA) at 200 µL/min. The cell was incubated for 10 min to bind the digoxygenin-labelled end of DNA to the cover glass surface. The unbound beads were removed by flushing the flow cell with 500 µL of MB at 200 µL/min.

Each of the two arrays (#Telo-18# and 601-20) was treated identically during the MMT experiments. Three parallel series of measurements were carried out for each of the three HO:DNA ratios applied in the two reconstitution runs using identical conditions for each array type.

Data collection and analysis were performed using procedures and scripts in the LabVIEW environment (National Instruments, USA) described in detail in^25, 60–63^. After manually adjusting the objective position, magnetic beads were automatically picked up and, if necessary, manually filtered, removing fault hits and double and stuck beads. Typically, 10 – 0 – 10 mm magnet shift of one or two 80-120 sec cycles of the fibres’ stretch – relief were recorded. The beads’ positions in three dimensions were monitored, applying recent 2D Fast Fourier Transforms algorithms to compute cross-correlations with computer-generated reference images^63^.

A statistical mechanics model was used to interpret the force-extension data of the nucleosome array stretching^60–62^. The model describes the dependence of total extension (*ztot*) of the nucleosome fibre along the direction of the applied force, *F*, (Extended Data Fig. 3e) as a sum of five stretching terms attributed to:

1). free DNA.
2). the folded chromatin fibre.
3). a beads-on-a-string chain after breaking nucleosome-nucleosome contacts and partial unwinding of an outer-turn DNA from the HO.
4). a beads-on-a-string chain consisting of HO’s or tetrasomes and hexasomes with a single wrap of 80 bp.
5). Fully unwrapped HO’s; continued stretching of the DNA results in largely irreversible histone loss.

Experimental force-extension curves were fitted to the model as described in detail in^25, 60^. First, experimental data were manually shifted to correspond to the length of fully stretched DNA, modifying z-coordinate offset and drift. Next, the unfolding model was fitted using a selection of fixed and variable parameters. The fixed parameters are either known, or expected numbers specific for a given nucleosome array (contour length of the template DNA, nucleosome repeat length, NRL), well-established properties of the free DNA (persistence length, stiffness), or parameters verified earlier in other studies, including tweezers measurements of the nucleosome arrays^25, 60, 64^.

The following values were explicitly fitted for each stretching curve:

i) The total number of nucleosome particles (nucleosomes, hexasomes, and tetrasomes) in the fibre, *N_total_*.
ii) The number of the nucleosome particles not included in the folded fibre, *N_unfold_*
iii) The fibre stiffness, *k_fibre_* (Extended Data Fig. 3e, stage 1).
iv) The free energy *ΔG_1_* is associated with the combined event of nucleosome unstacking and partial DNA unwrapping (Extended Data Fig. 3e, stage 2).
(5) *ΔG2*, free energy related to the second stage of DNA unwrapping up to a single turn of DNA (Extended Data Fig. 3e, stage 3).

The values of *N_total_* and *N_unfold_* were set manually by adjusting the fitting curve to the experimental points. *N_total_* was determined from the high-force part (> 8 pN) of the stretching curve corresponding to the ruptures of the last 79 bp DNA from the histone core; *N_unfold_* was set by fitting experimental data at the low-force range (0.5 – 2 pN). These unfolded nucleosomes undergo only the last-turn rupture event. The number of nucleosomes contributing to the fibre folding is defined as *Nfolded* = *N_total_* – *N_unfold_*. It is not possible^64^ to resolve the differences in the last rupture event between nucleosomes and tetrasomes, and it is assumed that both the populations have the same last step in the unwrapping pathway. The *k*, *ΔG_1_*, and *ΔG2* values were automatically fitted after initial guesses based on previous data. Data points obtained at forces larger than 7 pN were attributed to WLC curves with increasing lengths in steps of 79 bp.

The last stage of the nucleosome array stretching, largely irreversible ruptures of 75-80 bp DNA, was separately analysed using a dynamic spectroscopy equation modified to include a known number of the nucleosomes in the array at each of the rupture events^26, 65, 66^. Heat maps (Extended Data Fig. 3i-j) showing the dependence of rupture force events, *Frupt*, on the rate of applied force and number of nucleosomes in the array at the moment of rupture according to equation^26^:

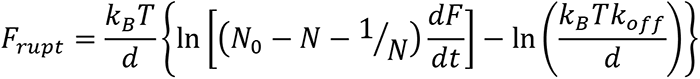

where *Frupt* is the rupture force; *d* is the distance between the bound state and the activation barrier peak along the direction of the applied force; N0 and N are the numbers of the nucleosomes at the initial moment and at the moment of rupture, respectively; *dF/dt* is the loading rate; *koff* is the rate constant for bond disruption under zero external force. Values of the *d* and *koff* were determined from the slope and intercept of the linear fit in coordinates *Frupt* versus 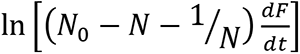.

## Reporting summary

### Data availability

Cryo-EM density maps for mono-NCP (EMD-31806), Di-NCP (EMD-31810 (3.9Å), EMD-31908 (5Å), EMD-31907 (4.6Å)), Open Di-NCP (EMD-31815(4.5 Å), EMD-31909 (6.6 Å), Tri-NCP (EMD-31816), Open Tri-NCP (EMD-31826) and Open Tetra NCP (EMD-31832), Telo-tetra (EMD-31823) and fitted models for mono-NCP (7V90), Di-NCP(7V96), Open Di-NCP(7V9C), Tri-NCP(7V9J), Open Tri-NCP (7V9S) and Open Tetra NCP (7VA4), Telo-tetra (7V9K) have been deposited in the Electron Microscopy Data Bank (EMDB) and the Protein Data Bank (PDB).

## Acknowledgements

We thank the NTU Institute of Structural Biology (NISB), Nanyang Technological University, for providing access to facilities. We acknowledge the Facility for Analysis, Characterisation, Testing and Simulation, Nanyang Technological University, for the use of their electron microscopy and Andrew Wong and Emmanuel Smith for assistance with Data collection. The authors acknowledge the Cryo-Electron Microscopy Facility at the Center for Bioimaging Science, Department of Biological Science, National University of Singapore, and Jian Shi for scientific and technical assistance. We are grateful to Curt Davey and Sara Sandin for valuable discussions and suggestions. This work has been supported by the Singapore Ministry of Education (MOE) Academic Research Fund (AcRF) Tier 2 [MOE2018-T2-1-112] and Tier 3 [MOE2012-T3-1-001] grants.

## Author contributions

AS, SYW, DR, and LN conceptualized and designed the study. AS prepared grids, collected and processed the Cryo-EM data, carried out model building and refinement, prepared constructs and biochemical assays. SYW prepared constructs and carried out negative stain EM, AUC experiments. SYW, WS, and SL designed and performed cloning and construct preparations. SL conceptualized and designed the MCS. SYW, WS, and QC prepared samples for force spectroscopy. VKV prepared grids and collected Cryo-EM data. NVB and JvN built the magnetic tweezer and performed measurements. NK and JvN performed and analysed force spectroscopy measurements. AS, WSY, DR, and LN wrote the paper with the help and input of all authors. All authors contributed to the interpretation of the data.

