## Supplementary Extended Data Figures 1-9 and tables 1-2 for "The columnar structure of human telomeric chromatin suggests mechanisms for telomere maintenance"

**This file includes:**

Extended Figures.1 to 9

Extended Tables 1 and 2

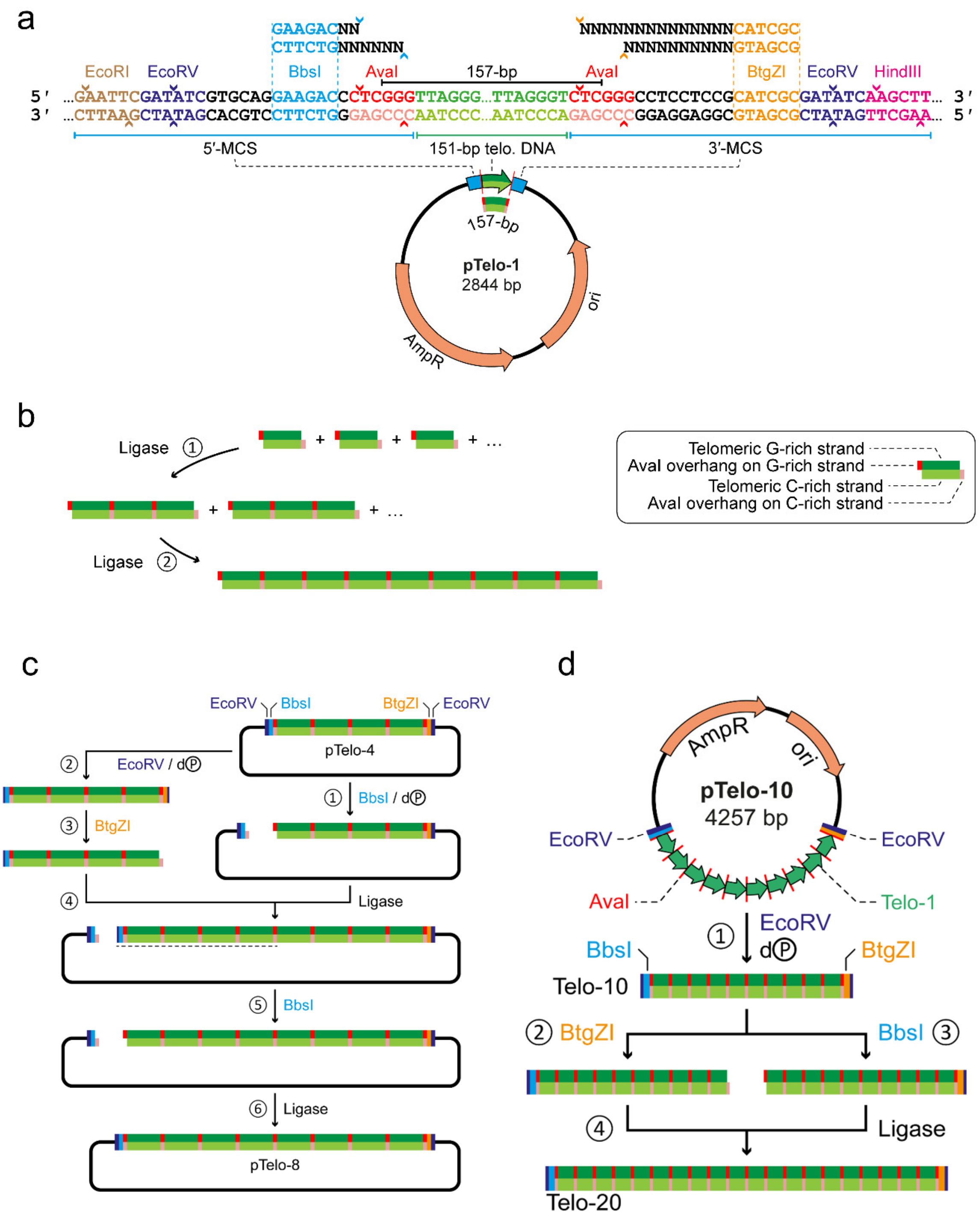


**Extended Data Fig. 1. Features and construction design of the pTelo constructs**. **a**, Plasmid map of the 2844 bp pTelo-1 construct. The pTelo-1 vector consists of a 151 bp human telomeric DNA insert flanked by two MCS (5′-MCS and 3′-MCS). The various restriction sites embedded inside the MCSs enable various manipulations and modifications of the telomeric insert.

Digestion with EcoRI and HindIII releases the telomeric insert bearing overhangs that can be ligated to asymmetric handles, such as those used for MMT measurements in this study. The two EcoRV sites enable releasing the telomeric insert with blunt ends for nucleosome reconstitution and the production of longer arrays. BbsI and BtgZI are Type IIS restriction enzymes that cleave the construct within the 5′- and 3′-AvaI sites, respectively. They let inserting additional array elements unilaterally or release telomeric arrays bearing AvaI overhangs suitable for multimerization of telomeric arrays. **b**, The elementary unit of the telomeric DNA template used in this study consists of a 151 bp human telomeric fragment (green) flanked by AvaI restriction sites (red) at each extremity. Once multimerized, the resulting arrays consist of integral multiples of 157 bp units. The 157 bp telomeric units are released from the pTelo-1 construct upon AvaI digestion and subsequently multimerized (1) into an array by ligation. The non-palindromic nature of the AvaI overhangs ensures control over strand orientation and prevents the G-rich and C-rich strands from mingling. The resulting multimers consist of integral multiples of 157 bp units, and the required multimer size is obtained by colony screening coupled with restriction digest analysis. (2) The entire process can be accelerated by ligating multimers together (e.g., 3× 157 bp repeat fragments into 6×, 9×, or 12× repeats). The release of 157 bp telomeric arrays bearing AvaI overhangs at each extremity is achieved by cutting the plasmid with BbsI and BtgZI. The resulting insert is subsequently ligated into integral multimers of the parental array. **c**, Alternatively, a pre-existing array can be extended unilaterally by selectively opening the vector either at the upstream or downstream AvaI site and inserting an extra telomeric array. This approach enables the array to be extended indefinitely in a controlled and stepwise process. Similarly, hybrid arrays bearing non-telomeric terminal sequences (e.g., 601 Widom or alpha-satellite DNA sequences) can be obtained using the same strategy. The array extension procedure is outlined in the Methods section and herein is illustrated with pTelo-4 plasmid. (1) Firstly, pTelo-4 is treated with BbsI and alkaline phosphatase to generate a linearized and dephosphorylated recipient vector with AvaI overhangs. (2) Separately, a Telo-4 array is excised from pTelo-4 by EcoRV digestion. (3) The resulting purified Telo-4 DNA fragment is digested with BtgZI to create a distal AvaI overhang and (4) ligated to the BbsI-linearized pTelo-4 from step 1. (5) The ligation product is subsequently deprotected with BbsI yielding a proximal AvaI overhang which enables the circularization of the pTelo-8 plasmid. **d**, Large-scale preparation of long telomeric DNA array template. (1) A 10×157-bp telomeric DNA array template (Telo-10) is excised from pTelo-10 vector by EcoRV digestion. The resulting purified DNA fragment is digested with BtgZI (2) or BbsI (3) to generate complementary AvaI overhangs at the downstream and upstream extremities. Subsequently, the resulting reaction products are ligated to yield a 20×157 bp telomeric DNA array template (Telo-20).

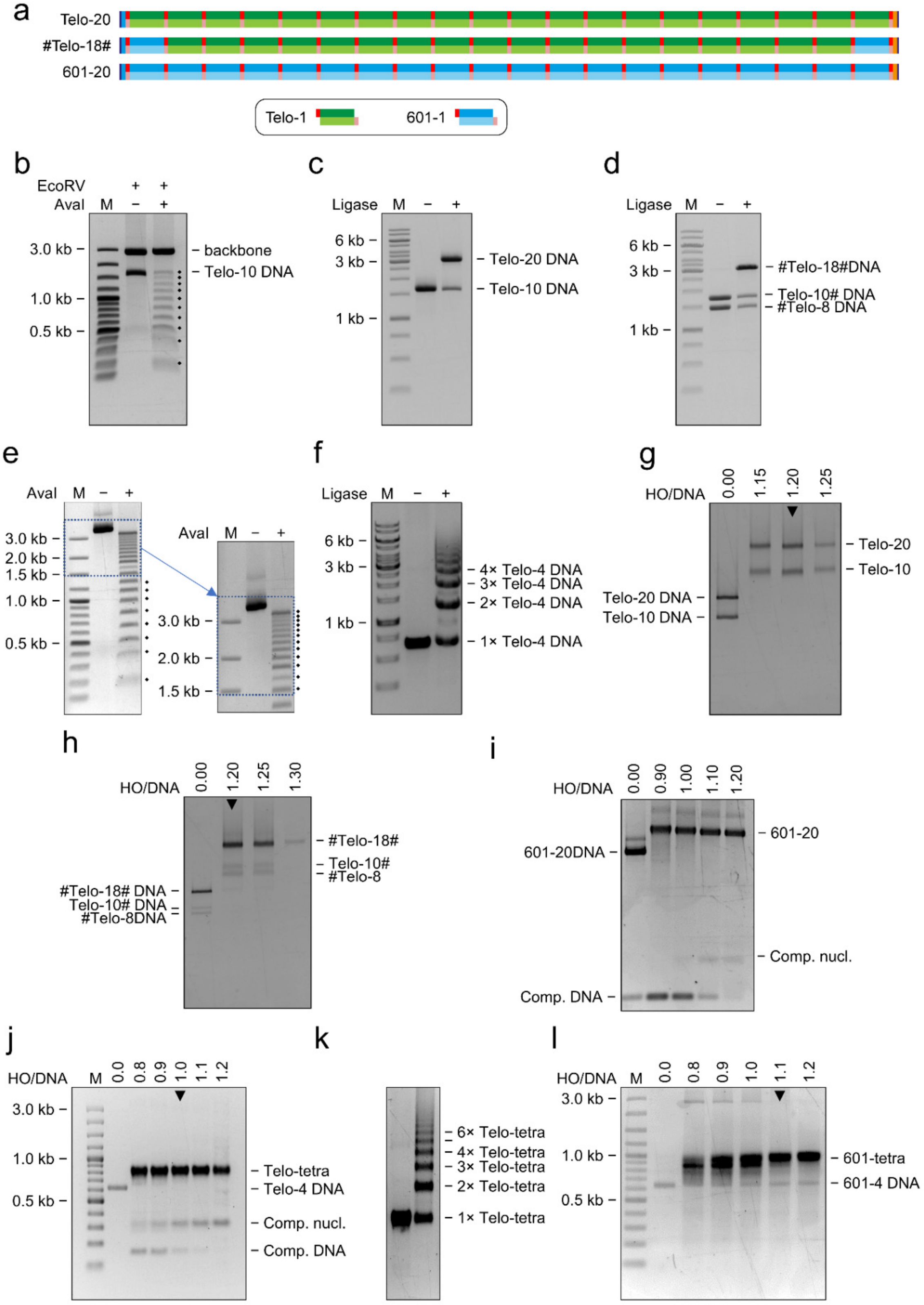


**Extended Data Fig. 2**. See next page for the caption.

**Extended Data Fig. 2. Preparation of Telomeric DNA and 601-20 DNA for in-vitro nucleosome array reconstitution. a**, Schematic diagram of various 20-mer 157 bp DNA array templates employed in this study. Telo-20 consists of twenty repeats of 157 bp Telo-1 fragment. #Telo-18# denotes a chimeric array template consisting of eighteen repeats of 157 bp Telo-1 unit flanked by one unit of 157 bp Widom 601 sequence (blue) at each terminus. 601-20 represents an array template made of twenty repeats of the 157 bp Widom 601 sequence. **b**, pTelo-10 was digested with EcoRV and AvaI for quality analysis. The resulting fragments were fractionated on 1.0% TBE-agarose gel. The quality of the amplified Telo-10 was assessed with EcoRV digestion of the plasmid to reveal any possible recombined telomeric DNA (lane 2). Additional partial digestion with AvaI (lane 3) showed the actual number of telomeric repeats the plasmid contains. **c**, An equimolar mixture of the BtgZI- and BbsI-digested Telo-10 fragments were ligated overnight and analyzed on 0.7% TBE-agarose gel. Densitometry analysis showed that the ligated mixture contained 70-75% of the Telo-20 DNA. **d**, Equimolar of BtgZI-digested #Telo-8 and BbsI-digested Telo-10# fragments were ligated overnight and analyzed on 0.7% TBE-agarose gel. Densitometry analysis showed that the ligated mixture contained 70-75% of the #Telo-18# DNA. **e**, Purified 601-20 DNA partially digested with AvaI to reveal exactly 20 repeats analyzed on 0.7% TBE-agarose gel. **f**, Telo-4 DNA template was released from the vector by EcoRV digestion followed by PEG size fractionation and ion-exchange purification (lane 2). This template was used to generate a DNA template containing multiple of Telo-4 by self-ligation mediated by T4 ligase and assisted by T4 polynucleotide kinase (lane 3) **g**, Telo-20 and **h** #Telo-18# reconstituted with recombinant human histone octamer titration analyzed on 0.7% Tris-borate agarose gel and post-stained with SYBR® Gold Nucleic Acid Gel Stain. **i**, 601-20 reconstituted with recombinant human histone octamer titration, with 147 bp pUC57 backbone fragment as competitor DNA. Reconstituted arrays were analyzed on 0.7% Tris-borate agarose gel and post-stained with SYBR® Gold Nucleic Acid Gel Stain. **j**, Telo-tetra was reconstituted with recombinant human histone octamer in the presence of 145 bp telomeric mononucleosomal DNA as competitor DNA to determine optimal saturation. The saturation of the reconstituted arrays was assessed by its migration on a 1.2% 0.25xTBE-agarose gel, and lane 1.0 indicates the ratio of optimal saturation. **k**, Reconstitution of self-ligated Telo-4 template to give arrays containing multiples of the telo-tetra unit. **l**, The 601-tetra nucleosome was reconstituted as a control for comparison with Telo-tetra. Lane 1.1 indicates the ratio with optimal saturation.

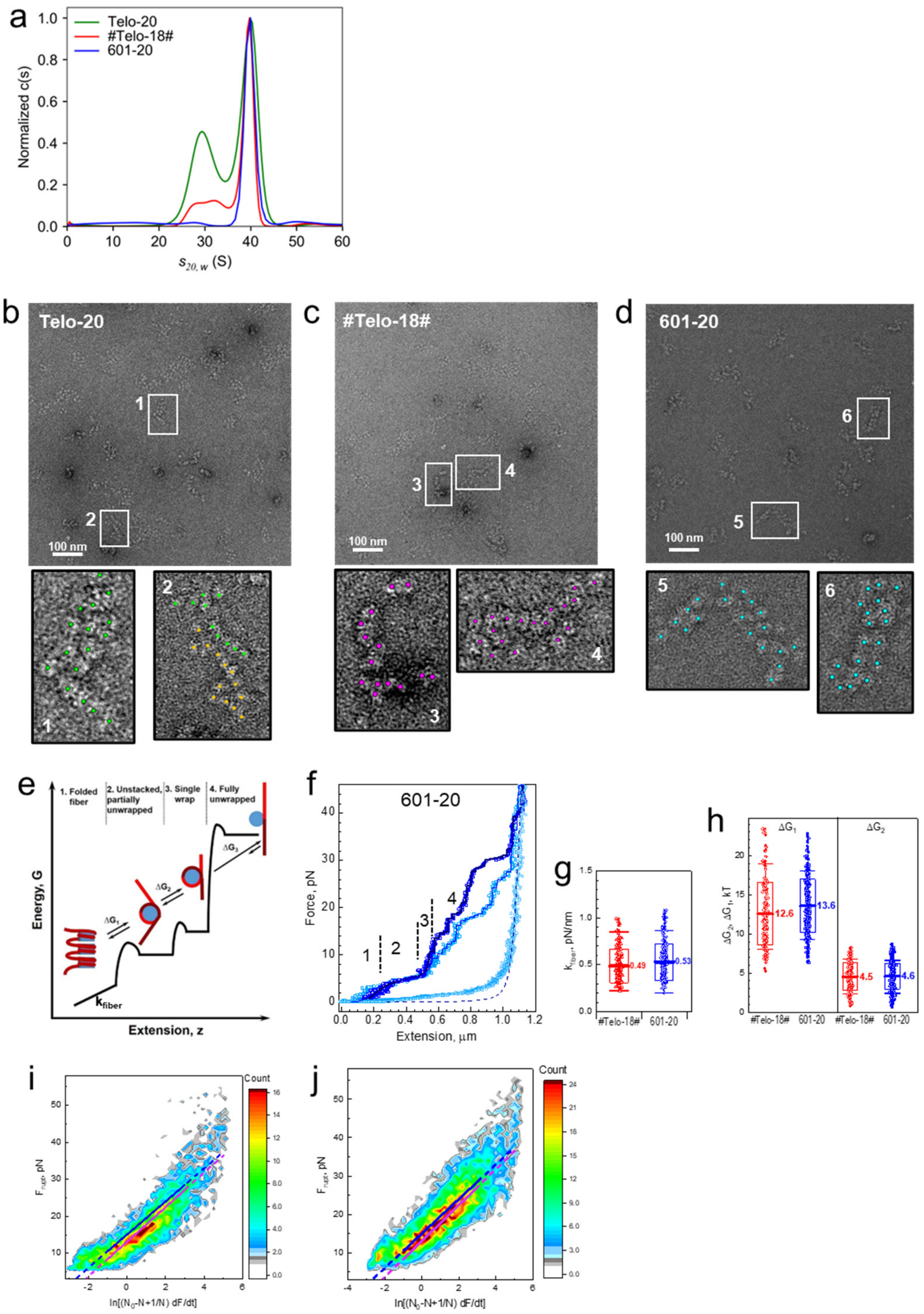


**Extended Data Fig. 3. AUC, EM and MMT analysis of the nucleosome arrays. a**, AUC-SV data normalized to maximum c(s) shows heterogeneity of the arrays. The s-value distribution of Telo-20 (green) shows 2 broad peaks at 39.4 S and 29.6 S, indicating the presence of respectively the long Telo-20 and the unligated Telo-10 arrays. The broad main peak of Telo-20 reflects the heterogeneity of the arrays caused by the lack of nucleosome position information in the telomeric sequence and the variation in nucleosome occupancy between individual fibres by ensemble-averaged measurements on fibres in solution unaffected by fixation. #Telo-18# (red) shows 3 overlapping peaks with s-values of 39.2 S, 32.5 S, and 27.8 S that reflect the presence of respectively #Telo-18#, unligated Telo-10# and #Telo-8 arrays. The sharper main peak of #Telo-18# distribution indicates lesser heterogeneity comparing to Telo-20. 601-20 (blue) shows a single peak at 40.2 S, demonstrates a homogenous array distribution. Representative negative stained electron-microscopy micrograph of **b** Telo-20 in 0.6 mM $Mg^{2+}$, **c** #Telo-18# in 0.6 mM $Mg^{2+}$, and **d** 601-20 in 0.8 mM $Mg^{2+}$. Boxes below each micrograph show blow-up of the selected single arrays chosen to illustrate stacked features of the telomeric arrays and ladder arrangement of the 601-20 array. (**e**) Free energy – extension scheme illustrating the statistical mechanics model of nucleosome array stretching. Fiber states are indicated at the top; four stages of fiber stretching are explained in the main text. (**f**) Examples of experimental force-extension curves recorded for the 601-20 array. The graph shows two recorded traces. Numbers refer to respective transitions shown in (**e**). (**g**) Fitted fiber stiffness, characterised by the stretching modulus ($k_{fiber}$) (**h**) Fitted free energies $\Delta G_1$ and $\Delta G_2$. In (**g**) and (**h**) data calculated for each trace are shown as small symbols; mean values are indicated in the graphs, boxes show standard deviation $\sigma$; error bars are for $2\sigma$. (**i**) and (**j**). Telomeric nucleosomes unwrap at lower forces than 601 nucleosomes. Heat maps showing the dependence of rupture force events, $F_{rupt}$, on the rate of applied force and the values of the *d* (the distance between the bound state and the activation barrier peak) and $k_{off}$ (the rate constant for bond disruption under zero external force) were determined as described in the Methods section. (**i**) and (**j**) are respectively the data for the #Telo18# and 601-20 arrays. Magenta (#Telo-18#) and blue (601-20) lines are linear fitting of the $F_{rupt}$ in the 10 pN < $F_{rupt}$ < 30 pN range. For the hybrid #Telo-18#, we obtained $d = 0.880 \pm 0.036$ nm and $k_{off} = 0.0128 \pm 0.0016$ 1/s; for the 601-20, - $d = 0.919 \pm 0.024$ nm and $k_{off} = 0.0077 \pm 0.0007$ 1/s.

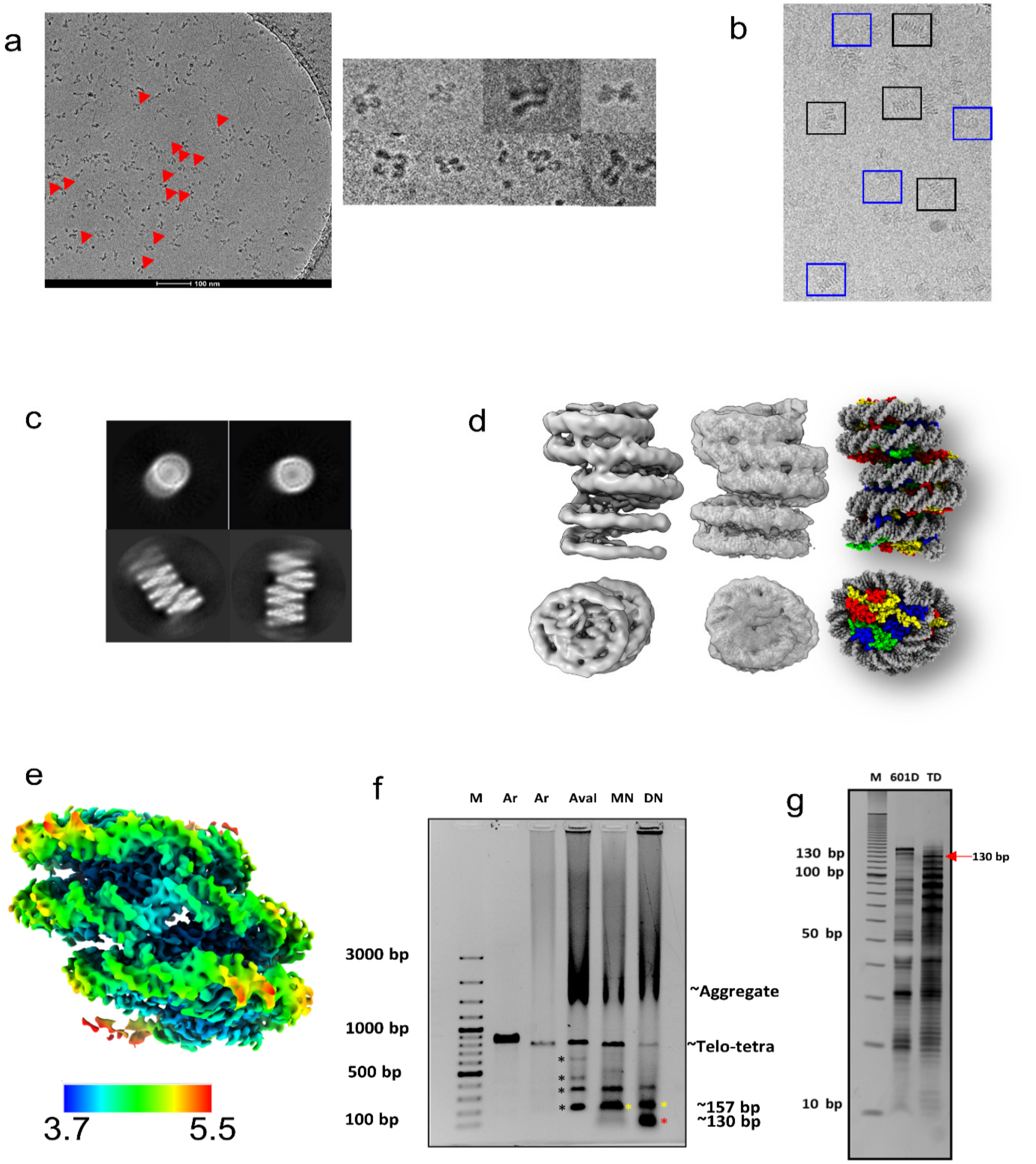


**Extended Data Fig. 4. 601-20 and Telo-tetra nucleosome arrays. a**, Sample Cryo-EM images of reconstituted 601-157 tetranucleosome showing zig-zag compaction under identical conditions at which telomeric tetranucleosome shows columnar packaging. Particles that exhibit the zig-zag arrangement are highlighted with a red arrow, and a close-up view is shown to the right. **b**, Representative micrograph of Telo-tetra arrays. Representative particles are shown in blue and black boxes, illustrating respectively a top view and side view. **c**, Representative 2Dclasses of Telo-tetra showing the top view and side views. **d**, The refined map of the Tri-NCP showed resolved features with minor densities for minor-major groves and helices of histones. The side and top view of the EM map, fitting and final structure is shown. **(e)** Local resolution estimation of Di-NCP (side view). **f**, AvaI, MNase, and DNase digestion of the Telo-tetra array. Lane M is a DNA marker; the two subsequent lanes marked Ar shows Telo-tetra arrays. Lane AvaI is an AvaI digest of the 4x157 array; lane MN is MNase digestion of the Telo-tetra array; lane marked DN is DNase digestion of the 4x157 array. The bands marked by a black asterisk in lane AvaI denote 1x157, 2x157, 3x157, and Telo-tetra bands are coming from AvaI digestion of the Telo-tetra array. MNase and DNase digest the array to nucleosome core length (145-157 bp) as observed by the DNA band marked by a yellow asterisk in lanes 5 and 6. They also digest into the nucleosome core as seen by the smear in lane MN and the DNA band marked by a red asterisk in lane DN. We have noted that under standard nuclease buffer, due to short linker length and presence of divalent ions significant amount of arrays precipitate to the bottom and is resistant to digestion. **g**, DNase digestion of telomeric and 601 NCP. We investigated the lack of 145 bp pause employing telomeric monoNCP. Lane M is a DNA marker, 601D shows DNase digestion of 601-NCP, TD shows DNase digestion of Telo-NCP. The 601 NCP shows a pause at 145 bp; however, the telomeric mononucleosome is readily digested with a pause at 130 bp.

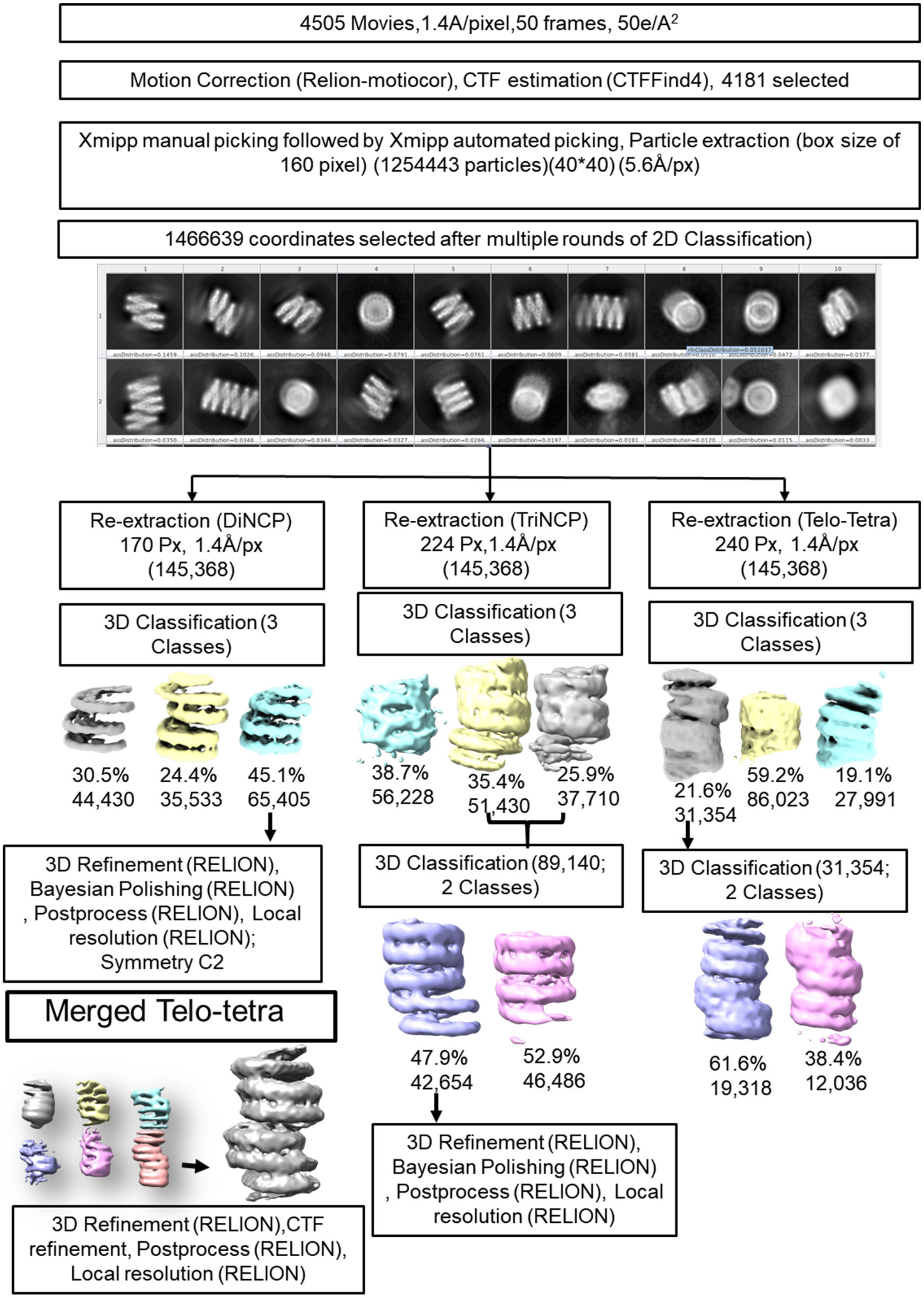


**Extended Data Fig. 5.** See next page for the caption.

**Extended Data Fig. 5. Flowchart of data processing of the Telo-tetra Dataset 1.** The flowchart shows the processing of the Dataset 1 collected at 1.4 Å/px. A separate dataset (Dataset 2) was collected containing 5766 micrographs (1.4 Å/pixel, 40 frames, 50 e/$Å^2$ ). It was processed identical to shown above, with the difference that picking was carried out employing a 3D template of Di-NCP shown above as reference. The Di-NCP from this Dataset 2 was processed to a resolution of 4.5 Å (Extended Data Fig. 7 e and f). For the Telo-tetra map, the maps obtained from a single dataset showed missing angular views. Hence, selected micrographs (8349) from two datasets were merged, followed by RELION auto picking employing tetra map from dataset 1 as 3D reference. The 1076747 coordinates picked went through multiple rounds of 2D classification from which 234492 coordinates were selected that went through 3D classification (bottom left box) to obtain a homogenous set of particles (26,580) corresponding to Telo-tetra that showed improved angular views and better resolution (8.1 Å) than individual datasets.

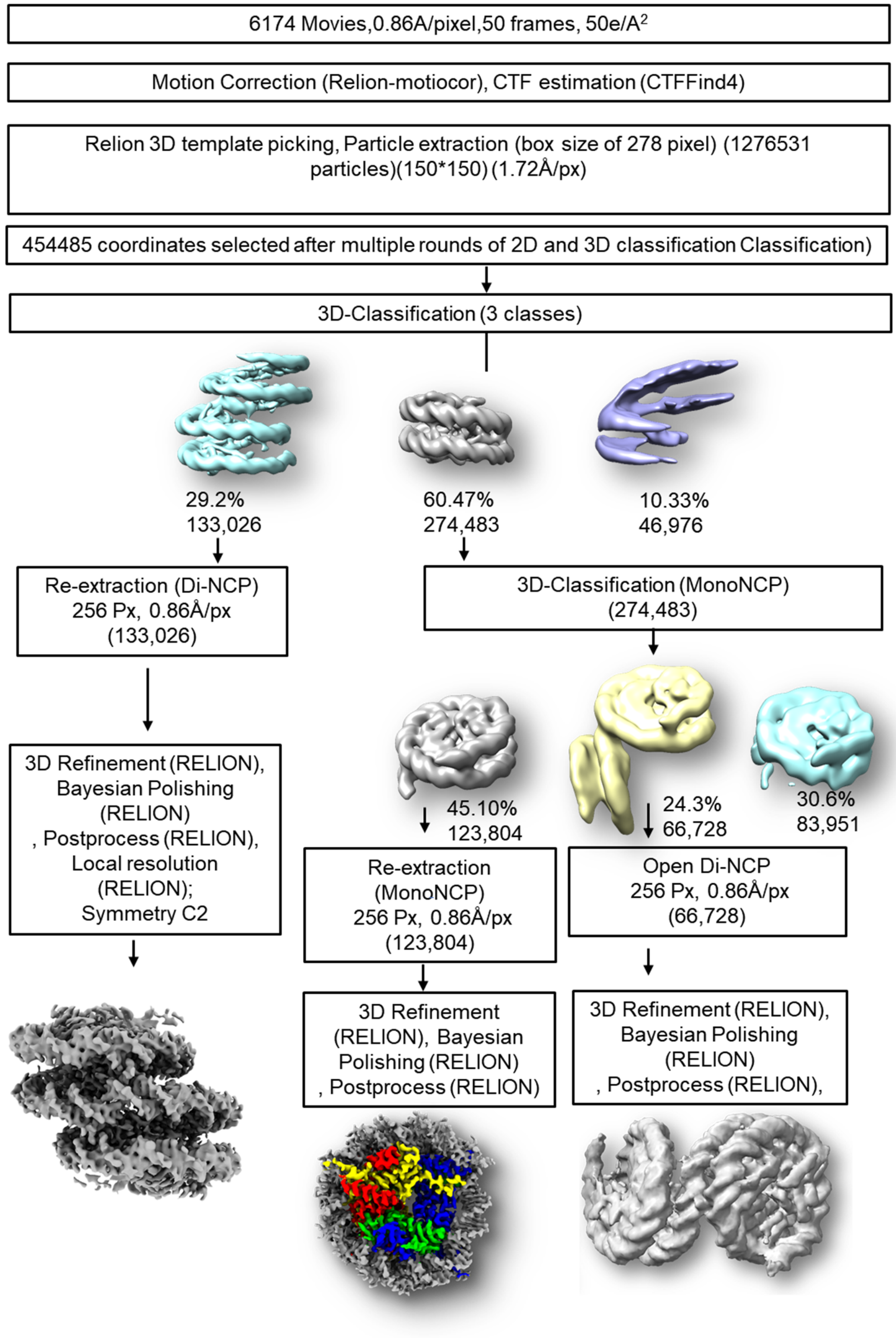


**Extended Data Fig. 6. Flowchart of data processing of the Telo-tetra Dataset 3, collected at 0.86 Å/pixel.** The processing of Dataset 3 was focused on the Di-NCP substructure of the Telo-tetra, employing the Di-NCP from Dataset 1 as a 3D template for picking. The higher-order structure from Dataset 3 did not show improvement in comparison to the merged higher-order structures and hence was not processed.

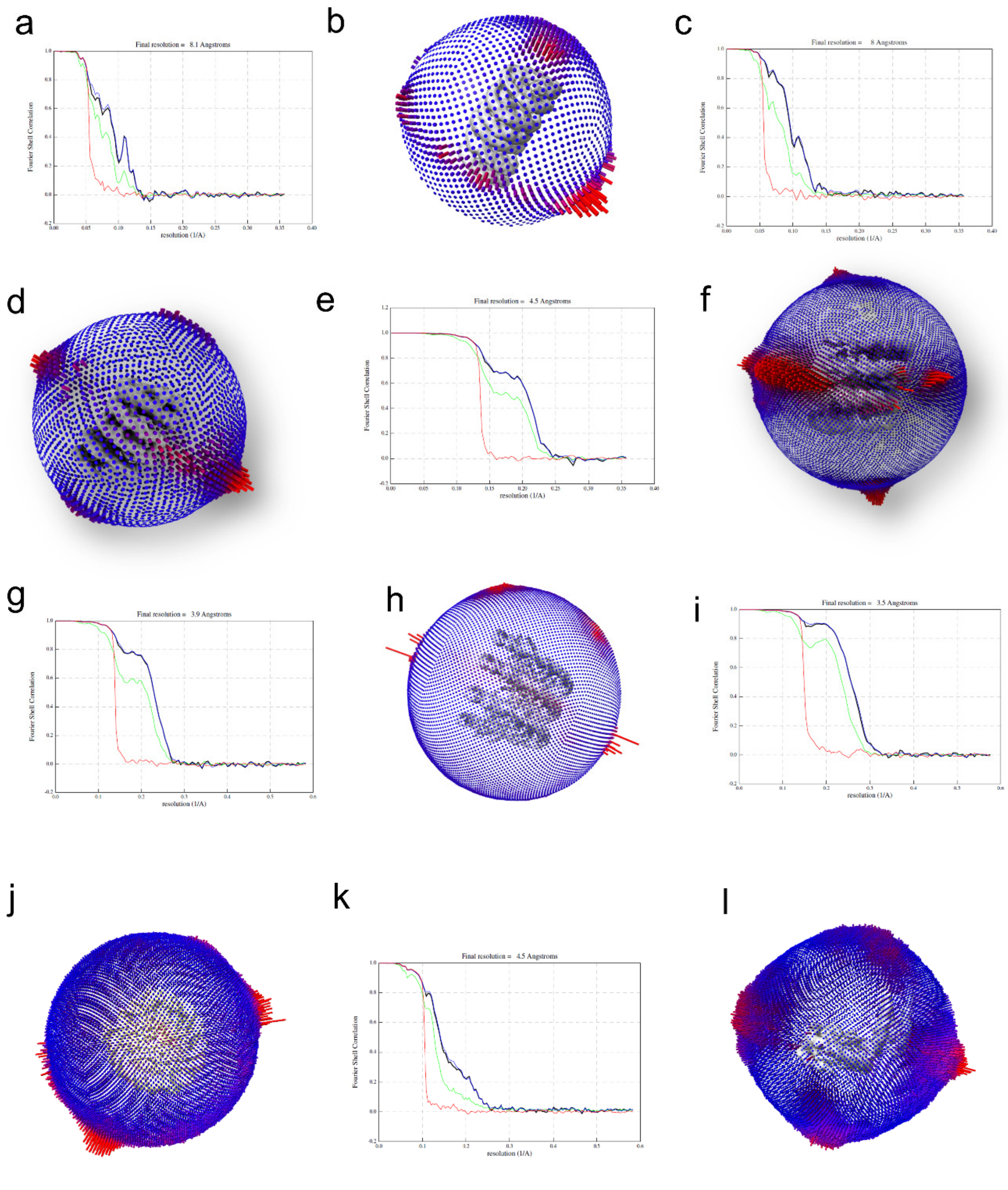


**Extended Data Fig. 7. Resolution and angular distribution of the Telo-tetra, Tri-NCP, and Di-NCP. a,** The Telo-tetra map was refined to a resolution of 8.1 Å. **b**, Angular distribution of particles used for refinement shows missing view and orientation preference. **c**, Tri-NCP volume resolution was estimated to a resolution of 8 Å. (**d**) Angular distribution of particles used for the Tri-NCP volume. **e**, The Di-NCP in 1.4 Å/px Dataset 2 was resolved to a resolution of 4.5 Å. **f**, Angular distribution of views of particles used for refinement. **g**, The Dataset 3 at 0.86 Å/pixel gave a resolution of 3.9Å for the Di-NCP. **h**, The angular distribution of particles used for the 3.9 Å Di-NCP map. **i**, The open Mono-NCP was resolved to a resolution of 3.5 Å. **j**, The angular distribution of Mono-NCP. **k**, The open Di-NCP was resolved to a resolution of 4.5 Å. **l**, The angular distribution of the open Di-NCP.

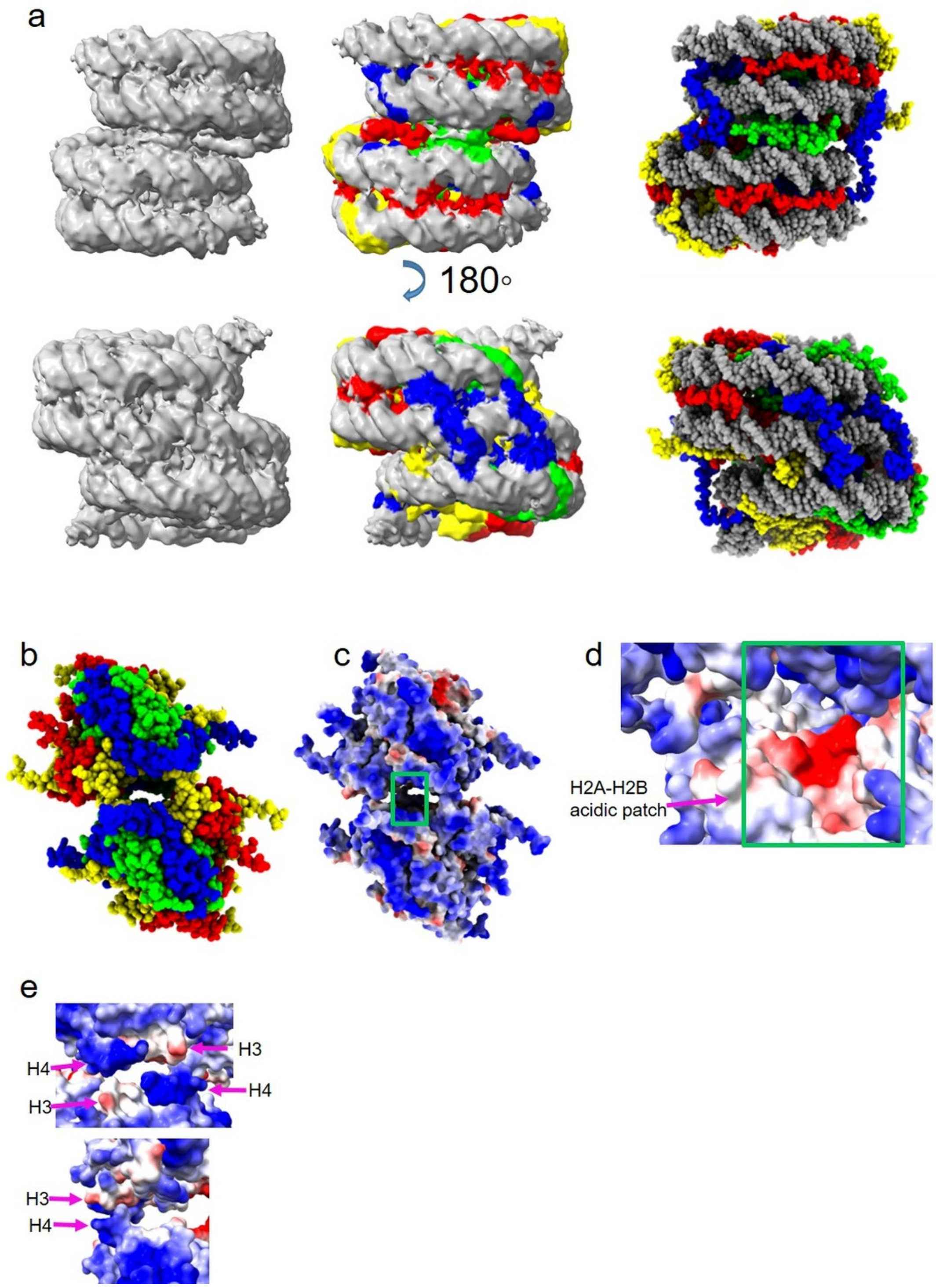


**Extended Data Fig. 8. Contribution of histone tails to Di-NCP stacking and substructures.**
**a**, EM densities arising from histone tail interaction with DNA were visible between the DNA gyres and along the minor groove (left panel). These bridging densities were assigned to individual histones (centre panel) based on the proximity of the EM density to the exit point of individual histone tails on the HO core. Multiple tube-like densities indicative of dynamic tails

were visible. The right panel shows the suggested location of all the histone tails as inferred from these tube-like fuzzy densities. The tube-like densities suggest multiple conformations for the tail, the most populated location of each tail are shown in the right panels. **b**, The histone octamer core depicting the H3-H2A clamp with histones H3, H4, H2A, and H2B coloured blue, green, yellow, and red. **c**, The histone octamer in panel a is shown with electrostatic surface colouring. The green box highlights the location of the H2A-H2B acidic patch, which is buried between the stacked histone cores. **d**, The H2A-H2B acidic patch, buried (green box in **c**) within the stacked octamers, does not interact with the histone core. **e**, The proximity of the two H4 tails (residues 21-24) to a weak acidic patch on the two H3 (residues 77-81) suggests two pairs of putative polar interaction that can supplement the H4 tail interaction with DNA (Fig. 3i).

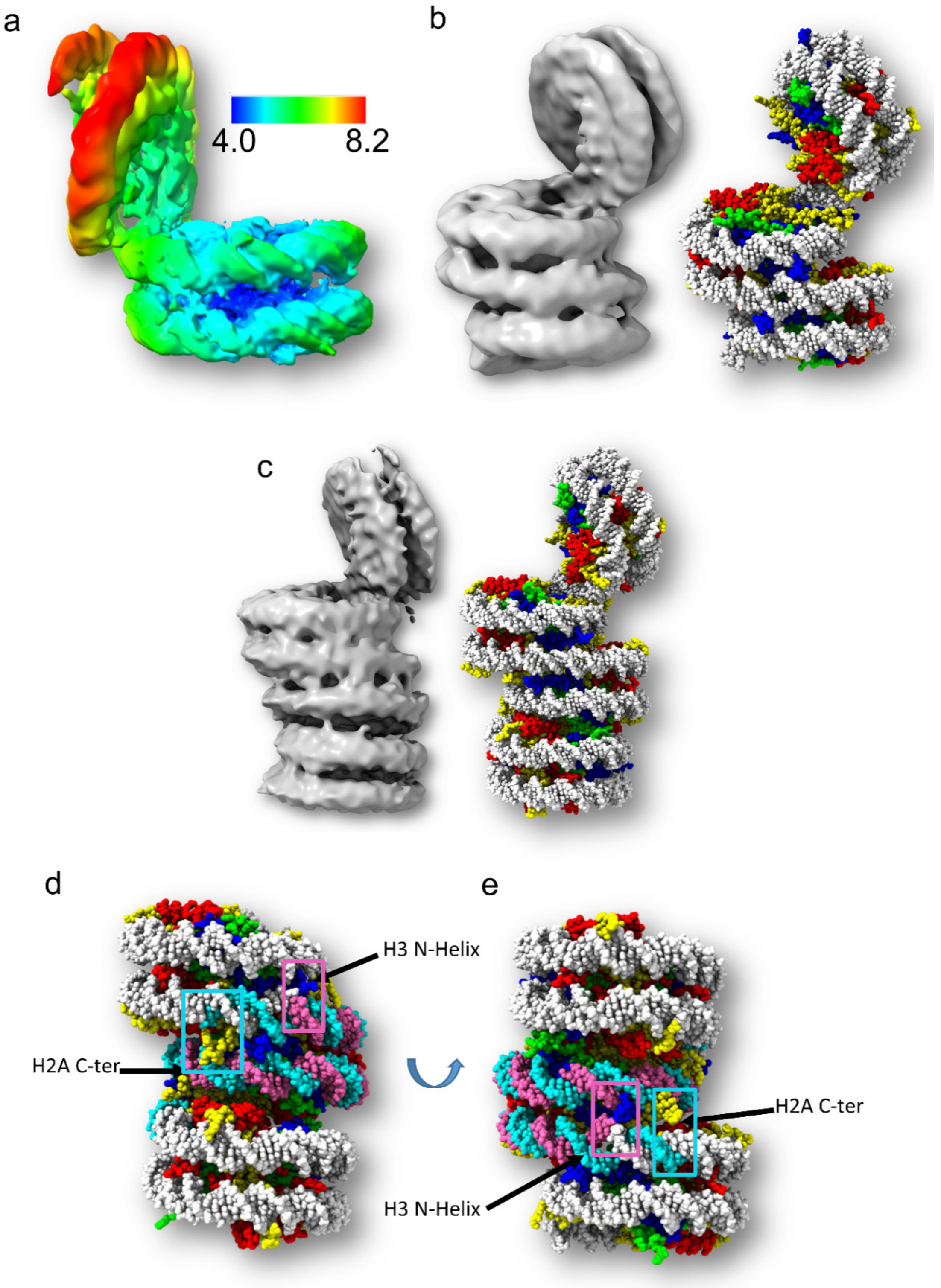


**Extended Data Fig. 9. Open and stacked conformations of the telomeric nucleosome arrays.**

**a**, Local resolution estimation of the open Di-NCP. **b,** EM map and structure of the open Tri-NCP. The map was reconstructed with 11,086 particles picked from dataset 2 at 1.4 Å/px. The particles used for the open Tri-NCP map comes from 130,951 coordinates that were discarded during Telo-tetra data processing. **c**, EM map and structure of open Telo-tetra. The map was reconstructed with 5,305 particles from dataset 2 selected from the 3D classification of the 130,951 particles. **d** and **e,** The structure of Telo-tetra results in the protection of DNA by the histones, specifically H2A (yellow sphere) and H3 (blue spheres). The H2A C-terminal protects DNA at the boundary of the histone octamers separated by ~157 bp (cyan spheres), suggesting the origin of the observed nuclease digestion band at 157 bp. Extended nuclease digestion results in digestion of the adjacent DNA positioned between the H2A C-terminal and the H3 helix, resulting in a ~130 bp band.

**Extended Data Table 1. Data collection and processing statistics of the Telo-tetra and sub-structures of Telo-tetra. Three datasets (Dataset 1, 2 and 3) were collected. All substructures were present in all three datasets, with minor variation in the population of the sub-structures between the datasets.**

| Sub-structure | Mono-NCP | Open Di-NCP | Di-NCP | Di-NCP | Di-NCP | Tri-NCP | Open Di-NCP | Open Tri-NCP | Open Telo-tetra | Telo- tetra |
|---|---|---|---|---|---|---|---|---|---|---|
| **Sample** | Telomeric tetranucleosome | | | | | | | | | |
| Sample support | Quantifoil R1.2/1.4 | | | | | | | | | |
| Magnification | 105000 | | | | | | | | | |
| Voltage (kV) | 300 | | | | | | | | | |
| Electron exposure (e–/Å$^2$) | 50 | | | | | | | | | |
| Defocus range (µm) | 0.5-2.5 | | | | | | | | | |
| **Detector** | K3 | | | K2 | | | | | | |
| Pixel size (Å) | 0.86 | | | 1.4 | | | | | | |
| Dataset origin | Dataset 3 | | | Dataset 1 | Dataset 2 | Dataset 1 | Dataset 2 and 1 | Dataset 2 | | Dataset 2 and 1 |
| Symmetry imposed | C1 | | C2 | | | C1 | | | | C2 |
| Initial particle images (no.) | 1,276,531 | | | 1,254,443 | 1,661,680 | 1,254,443 | 2,340,179 | 816,663 | | 1,076,747 |
| Final particle images (no.) | 123,804 | 82,175 | 133,026 | 63,501 | 136,537 | 42,654 | 62136 | 11,086 | 5305 | 26,580 |
| Map resolution (Å) FSC threshold (0.143) | 3.5 | 4.5 | 3.9 | 5 | 4.6 | 8 | 6.6 | 10.9 | 14.3 | 8.1 |
| EMDB | EMD-31806 | EMD-31815 | EMD-31810 | EMD-31908 | EMD-31907 | EMD-31816 | EMD-31826 | EMD-31832 | EMD-31909 | EMD-31823 |
| PDB | 7V90 | 7V9C | 7V96 | | | 7V9J | | 7V9S | 7VA4 | 7V9K |

### Extended Data Table 2. Structure refinement statistics of Telo-tetra and sub structures.

| EMDB map<br>PDB<br>Sub-structure | EMD-31806<br>PDB- 7V90<br>Mono | EMD-31815<br>PDB-7V9C<br>Open Di-NCP | EMD-31810<br>7V96<br>Di-NCP | EMD-31816<br>7V9J<br>Tri-NCP | EMD-31826<br>7V9S<br>Open Tri-NCP | EMD-31832<br>7VA4<br>Open Telo-tetra | EMD-31823<br>7V9K<br>Telo-tetra |
|---|---|---|---|---|---|---|---|
| Initial model used<br>(PDB code) | 6KE9 | | | | | | |
| Model resolution (Å)<br>FSC threshold<br>(0/0.143/0.5) | 3.0/3.1/3.6 | 3.46/4.27/8.01 | 3.75/3.84/4.06 | NA | NA | NA | NA |
| Model composition<br>Non-hydrogen atoms<br>Protein residues<br>Nucleotide | <br>12148<br>781<br>290 | <br>23228<br>1518<br>546 | <br>23968<br>1593<br>550 | <br>35276<br>2376<br>798 | <br>34909<br>2375<br>781 | 46944<br>3128<br>1078 | <br>47361<br>3173<br>1078 |
| *B* factors (Å$^2$)<br>Protein<br>DNA<br>(min/max/mean) | <br>0.00/164.31/36.92<br>20.00/214.81/135.18 | <br>0.50/702.40/57.72<br>20.00/999.00/308.71 | <br>0.50/30.00/1.83<br>0.50/20.00/0.73 | <br>0.00/164.31/13.37<br>20.00/244.28/177.00 | <br>0.00/164.31/13.36<br>0.00/164.31/13.36 | <br>0.50/30.00/1.44<br>0.50/20.00/0.95 | <br>0.00/164.31/10.47<br>0.50/20.00/0.75 |
| R.m.s. deviations<br>Bond lengths (Å)<br>Bond angles (°) | <br>0.005<br>0.851 | <br>0.003<br>0.663 | <br>0.011<br>1.620 | <br>0.004<br>0.696 | <br>0.007<br>0.944 | 0.004<br>0.711 | 0.004<br>0.707 |
| Validation<br>MolProbity score<br>Clashscore<br>Poor rotamers (%) | <br>2.27<br>8.41<br>4.01 | <br>1.94<br>8.56<br>1.27 | <br>2.64<br>6.37<br>15.66 | <br>2.16<br>8.77<br>1.97 | <br>2.31<br>4.33<br>7.03 | 2.47<br>10.59<br>2.66 | <br>2.20<br>11.61<br>1.52 |
| Ramachandran plot<br>Favored (%)<br>Allowed (%)<br>Disallowed (%) | <br>94.77<br>4.71<br>0.52 | <br>94.08<br>4.98<br>0.94 | <br>94.49<br>4.74<br>0.77 | <br>92.65<br>6.36<br>0.99 | <br>92.78<br>6.36<br>0.86 | 87.63<br>9.79<br>2.58 | <br>92.09<br>6.34<br>1.58 |
